# Horizontal transfer and recombination fuel Ty4 retrotransposon evolution in *Saccharomyces*

**DOI:** 10.1101/2023.12.20.572574

**Authors:** Jingxuan Chen, David J. Garfinkel, Casey M. Bergman

**Affiliations:** Institute of Bioinformatics, University of Georgia, Athens, GA, USA; Department of Biochemistry and Molecular Biology, University of Georgia, Athens, GA, USA; Department of Genetics, University of Georgia, Athens, GA, USA

**Keywords:** genome evolution, horizontal transfer, recombination, retrotransposon, yeast

## Abstract

Horizontal transposon transfer (HTT) plays an important role in the evolution of eukaryotic genomes, however the detailed evolutionary history and impact of most HTT events remain to be elucidated. To better understand the process of HTT in closely-related microbial eukaryotes, we studied Ty4 retrotransposon subfamily content and sequence evolution across the genus *Saccharomyces* using short- and long-read whole genome sequence data, including new PacBio genome assemblies for two *S. mikatae* strains. We find evidence for multiple independent HTT events introducing the Tsu4 subfamily into specific lineages of *S. paradoxus*, *S. cerevisiae*, *S. eubayanus*, *S. kudriavzevii* and the ancestor of the *S. mikatae*/*S. jurei* species pair. In both *S. mikatae* and *S. kudriavzevii* , we identified novel Ty4 clades that were independently generated through recombination between resident and horizontally-transferred subfamilies. Our results reveal that recurrent HTT and lineage-specific extinction events lead to a complex pattern of Ty4 subfamily content across the genus *Saccharomyces*. Moreover, our results demonstrate how HTT can lead to coexistence of related retrotransposon subfamilies in the same genome that can fuel evolution of new retrotransposon clades *via* recombination.

## Introduction

Transposable elements (TEs) are mobile, repetitive DNA sequences that are found in nearly all eukaryotic genomes. Typically, TEs are vertically inherited from parents to offspring within a species (Wells and Feschotte, 2020). However, transmission of TEs between different species can also occur through the process of horizontal transposon transfer (HTT) (Daniels *et al*., 1990). With the widespread availability of whole genome sequencing data, HTT has been increasingly recognized as an important phenomenon in the evolution of eukaryotic genomes (Schaack *et al*., 2010; Wallau *et al*., 2012; Gilbert and Feschotte, 2018; Wallau *et al*., 2018).

HTT events can be identified using different sources of evolutionary evidence including phylogenetic incongruence between host genomes and TE sequences, unexpectedly high sequence similarity between TEs from divergent species, or the patchy distribution of a TE family across a set of closely-related lineages (Wallau *et al*., 2012; Peccoud *et al*., 2018). Based on these criteria, an increasing number of HTT events have been identified across the tree of life, with many examples in plants and animals (Dotto *et al*., 2015). While the number of HTT events detected in fungi is still relatively rare (Dobinson *et al*., 1993; Daboussi *et al*., 2002; Novikova *et al*., 2009; Amyotte *et al*., 2012; Sarilar *et al*., 2015), a growing number of HTT events have been identified among *Saccharomyces*yeast species (Liti *et al*., 2005; Carr *et al*., 2012; Bergman, 2018; Czaja *et al*., 2020; Bleykasten-Grosshans *et al*., 2021). As in other taxa, previous studies reporting HTT in yeast have typically focused on detecting their existence. However, the timing, geographic location, identity of donor/recipient lineages, and consequences of these HTT events remain unknown.

The Ty4 long-terminal repeat (LTR) retrotransposon family is a promising model to understand the process and impact of HTT in *Saccharomyces* yeasts. Ty4 elements – like other members of the Ty1/Copia superfamily – are composed of two overlapping ORFs (*gag* and *pol*) flanked by LTRs (Janetzky and Lehle, 1992; Stucka *et al*., 1992). The founding member of the Ty4 family was first identified in *S. cerevisiae* (Stucka *et al*., 1989) and defined the eponymous Ty4 subfamily. After the initial discovery of the Ty4 subfamily, Neuveglise *et al*. (2002) described a related subfamily called Tsu4 in the distant congener *S. uvarum*. More recently, Bergman (2018) reported the unexpected presence of Tsu4 sequences in *S. paradoxus*, *S. cerevisiae*, and *S. mikatae* and proposed these observations could be explained by HTT events from a donor related to *S. uvarum* or its sister species *S. eubayanus*. The strongest evidence for HTT involving the Tsu4 subfamily was found in *S. paradoxus* based on its patchy distribution among divergent *S. paradoxus* lineages, a high similarity between *S. paradoxus* Tsu4 elements and those from *S. uvarum* and *S. eubayanus*, and discordance between the phylogeny of Tsu4 elements and the host tree of *Saccharomyces* species. Bergman (2018) also reported evidence for a potential Tsu4 HTT event involving *S. cerevisiae* based on the presence of one Tsu4 full-length element (FLE) in a single strain (called 245) and its high sequence similarity with the Tsu4 sequences recently introduced into *S. paradoxus*. Subsequently, O’Donnell *et al*. (2023) confirmed the presence of Tsu4 sequences in *S. cerevisiae* in a different strain (called CQS), although it is currently unclear whether Tsu4 in strains 245 and CQS arose from the same or different HTT events. Likewise, Bergman (2018) provided limited evidence for a possible third Tsu4 HTT event in *S. mikatae* based on discordance between the phylogeny of Tsu4 elements and the accepted tree for the genus *Saccharomyces* (Borneman and Pretorius, 2015).

Despite these advances, several important aspects of how HTT events affect the evolution of the Ty4 family in *Saccharomyces* yeasts remain unresolved. First, the samples of *S. cerevisiae* and *S. paradoxus* genomes previously studied did not span the global diversity of these species. Thus the timing, geographic origin, and number of Tsu4 HTT events in these species is not fully understood. Second, previous inferences about the potential donor species for the Tsu4 HTT event into *S. paradoxus* were limited by the lack of high-quality whole genome assemblies (WGAs) for *S. uvarum* and *S. eubayanus*. For example, the inference that Tsu4 elements from *S. eubayanus* are most closely related to those transferred in *S. paradoxus* (Bergman, 2018) was made indirectly using genome data from the interspecific hybrid species *S. pastorianus*, which contains subgenomes from *S. eubayanus* and *S. cerevisiae* (Libkind *et al*., 2011; Baker *et al*., 2015; Okuno *et al*., 2016). Third, evidence for the putative Tsu4 HTT in *S. mikatae* was based on a single element from a highly fragmented draft WGA (Cliften *et al*., 2003), which are known to have incompletely reconstructed TE sequences (Myers *et al*., 2000). Finally, the single Ty4 family member identified in a draft genome for *S. kudriavzevii* (which is a key outgroup species to *S. cerevisiae*, *S. paradoxus*, and *S. mikatae*) was found to be in an intermediate phylogenetic position to both the Ty4 and Tsu4 subfamilies (Bergman, 2018). These results suggest that additional subfamilies in the Ty4 family remain to be discovered, or that recombination occurred between the Ty4 and Tsu4 lineages. Here we use large-scale short-read resequencing data and high-quality long-read WGAs from multiple species in the genus *Saccharomyces* to address the impact of HTT on the evolution of the Ty4 family. Our results support a complex model for Ty4 family evolution in yeast that is shaped by recurrent HTT events involving the Tsu4 subfamily, lineage-specific extinction events, and creation of new retrotransposon clades through recombination between pre-existing subfamilies that co-occur in the same species because of HTT.

## Results and Discussion

To better understand the history and impact of HTT events involving the Ty4 family in *Saccharomyces*, we used four complementary genomic strategies. First, to more accurately infer the biogeographic distribution and ancestral states of the Ty4/Tsu4 subfamilies in *S. paradoxus* and *S. cerevisiae*, we investigated the presence/absence of Ty4/Tsu4 subfamily sequences in worldwide phylogenies for both species using unassembled short-read WGS datasets. For these analyses, we estimated copy number for LTRs and internal regions separately because recombination between LTRs within FLEs frequently excises internal sequences creating solo LTRs (Farabaugh and Fink, 1980). Doing this allows us to interpret the presence of internal sequences as evidence of recent activity, and the presence of LTRs as evidence of both recent and past activity in a strain. Second, we annotated Ty4/Tsu4 copies in a dataset of over 200 high-quality WGAs for all species in the *Saccharomyces* genus which allowed us to cross-validate Ty4/Tsu4 subfamily presence/absence data based on short-read WGS data, and to classify annotated copies at higher resolution into FLEs, truncated elements, and solo LTRs. We interpret the presence of FLEs in a WGA as evidence for recent activity in a strain, while truncated elements and solo LTRs represent past activity. Third, we generated phylogenetic networks and trees for internal coding regions of FLEs extracted from WGAs, which allowed us to directly investigate the molecular evolution of the Ty4 family across the entire *Saccharomyces* genus. Fourth, we generated strainspecific consensus sequences from unassembled short-read WGS datasets, which allowed us to study Tsu4 subfamily evolution among *S. paradoxus*, *S. cerevisiae*, *S. eubayanus* and *S. uvarum* using larger samples of strains that lack high-quality WGAs.

### An ancestral Tsu4 HTT event occurred prior to radiation of indigenous American *S. paradoxus* lineages

Using short-read WGS datasets for 370 *S. paradoxus* strains, we reconstructed a maximum likelihood (ML) phylogenetic tree based on 713,556 SNPs that confirmed all major known lineages, sub-lineages, and their relationships (Figure 1A) (Naumov *et al*., 1997; Koufopanou *et al*., 2006; Kuehne *et al*., 2007; Liti *et al*., 2009; Leducq *et al*., 2016; Henault *et al*., 2019; Eberlein *et al*., 2019; He *et al*., 2022). Worldwide diversity in *S. paradoxus* splits into the two major known lineages: American and Eurasian. The American lineage includes several indigenous North American sub-lineages (SpB, SpC, SpC* and SpD), as well a lineage with a single strain from Hawaii. SpC* and SpD are hybrid lineages derived from crosses between SpB and SpC, and between SpB and SpC*, respectively (Leducq *et al*., 2016; Henault *et al*., 2019; Eberlein *et al*., 2019). The Hawaiian lineage has been reported to share similarity with either the SpB (Leducq *et al*., 2014) or SpC/SpC* lineages (He *et al*., 2022; Peris *et al*., 2023). Our analysis places the Hawaiian lineage as an outgroup to the SpC/SpC* lineages (circled number 3, Figure 1A). Importantly, we note that *S. paradoxus* lineage from S. America (circled number 2, Figure 1A) – which was formerly considered a distinct species called *S. cariocanus* (Naumov *et al*., 2000) – is contained within the North American SpB sub-lineage (Koufopanou *et al*., 2006; Liti *et al*., 2006, 2009; Hyma and Fay, 2013; Leducq *et al*., 2014; He *et al*., 2022). The Eurasian lineage includes sub-lineages indigenous to Europe, Far East Asia, and China, as well as a sub-lineage (SpA) composed of strains from North America that descend from a recent trans-oceanic migration event (Kuehne *et al*., 2007; Leducq *et al*., 2016).

**Figure 1:**
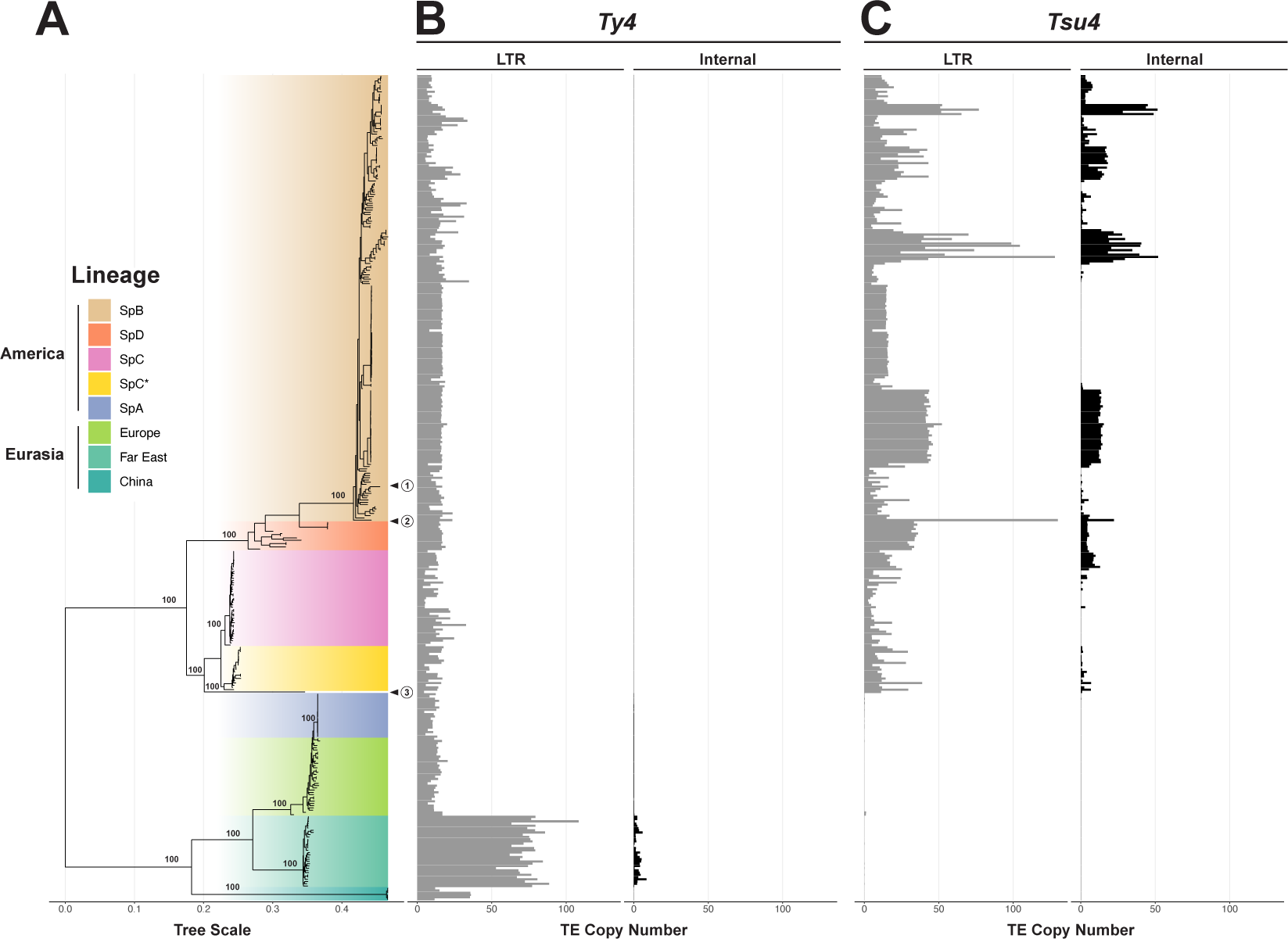
Host phylogeny of *S. paradoxus* annotated with estimated Ty4/Tsu4 copy number. (A) Midpoint rooted ML phylogenetic tree of 370 *S. paradoxus* strains integrated from multiple public short-read WGS datasets (see Materials and Methods). Bootstrap support is annotated for key nodes. Major lineages and sub-lineages are annotated according to previously-reported population structure (Eberlein *et al*., 2019; He *et al*., 2022). The N. American strain DG1768 used in retromobility studies (Chen *et al*., 2022b) is found in the SpB sub-lineage and is indicated by the circled number 1. The S. American strain UFRJ50916 is found in the SpB sub-lineage and is indicated by the circled number 2. The Hawaiian strain UWOPS91-917.1 is found in its own sub-lineage indicated by the circled number 3. (B) Copy number estimates for the Ty4 subfamily. (C) Copy number estimates for the Tsu4 subfamily. In (B) and (C), gray bars represent copy number estimates for LTRs, whereas black bars represent estimated copy number for internal regions.

By mapping estimated Ty4/Tsu4 subfamily copy number onto the global phylogeny for *S. paradoxus*, we observe that Ty4 LTR sequences are present in all *S. paradoxus* strains from both the Americas and Eurasia (Figure 1B). Strains from the Far East sub-lineage show a significantly higher Ty4 LTR copy number in comparison to other sub-lineages. Ty4 internal regions are essentially absent across the species, except in the LTR-rich Far East sub-lineage (Figure 1B). In contrast, Tsu4 LTR sequences are only found in indigenous American strains and absent from strains with a Eurasian origin (Figure 1C). Tsu4 internal sequences are found in all indigenous American sub-lineages (SpB, SpC, SpC*, SpD, and Hawaii) with highly variable copy number (Figure 1C). Notably, we found no Tsu4 sequences in the Eurasian-derived North American SpA sub-lineage (see also (Henault *et al*., 2020)), which shows no evidence of admixture with indigenous American sub-lineages after secondary contact (Kuehne *et al*., 2007; Hyma and Fay, 2013; Leducq *et al*., 2014, 2016; He *et al*., 2022).

Analysis of Ty4/Tsu4 subfamily content in Ty4 sequences in a smaller dataset of 12 long-read WGAs that samples all major *S. paradoxus* lineages cross-validated results based on short-read WGS data. Ty4 solo LTRs were found in all strains but Ty4 FLEs were only found in the Far East strain N44 (Table S1). In contrast, Tsu4 solo LTRs are identified in the nine indigenous American *S. paradoxus* strains, and are absent from the other three strains with Eurasian origin (CBS432, N44, and

LL2012_001) (Table S1). At least one Tsu4 FLE is identified in all indigenous American *S. paradoxus* strains with WGAs except for the SpB strain DG1768 that is commonly used in retromobility studies (Chen *et al*., 2022b) (circled number 1, Figure 1A). As previously reported (Bergman, 2018), Tsu4 copy number in the South American SpB strain UFRJ50916 is much higher than other *S. paradoxus* strains with WGAs.

These data indicate that the Ty4 subfamily was present in the most recent common ancestor (MRCA) of all *S. paradoxus* lineages prior to global dispersal, and therefore represents the ancestral subfamily in *S. paradoxus*. The Ty4 subfamily subsequently went extinct in most recognized *S. paradoxus* sub-lineages except for the Far East sub-lineage where it remains active. In contrast, the lack of Tsu4 sequences in Eurasian *S. paradoxus* and the Eurasian-derived SpA sub-lineage indicates this subfamily has never existed in Eurasia and therefore was not present in the MRCA of all *S. paradoxus* strains. Our results support the interpretation that a Tsu4 HTT event occurred in an ancestor of all indigenous American *S. paradoxus* sub-lineages after the divergence of American from Eurasian lineages. This HTT event most likely occurred in an ancestral lineage where the Ty4 subfamily had already gone extinct, thus explaining why Ty4 and Tsu4 FLEs have never been observed in the same *S. paradoxus* strain. Since this ancestral HTT event, Tsu4 has maintained activity in all indigenous American *S. paradoxus*sublineages. However, Tsu4 has secondarily gone extinct or expanded to very high copy-number in many strains in each American *S. paradoxus* sub-lineage. We note that this parsimonious scenario does not exclude the possibility of additional recent Tsu4 HTT events into indigenous American *S. paradoxus* lineages that are obscured by this initial ancestral HTT event.

### Recent HTT has introduced Tsu4 into a small number of Central/South American *S. cerevisiae* strains

Using a similar short-read WGS-based approach as was used for *S. paradoxus*, we reconstructed a species-wide phylogeny for *S. cerevisiae* based on 2,787,577 genome-wide SNPs from 2,404 strains (Figure 2). In contrast to the ML approach used for *S. paradoxus* where admixture among lineages is rare, we followed Peter *et al*. (2018) in using a neighbor-joining (NJ) approach to generate the *S. cerevisiae* phylogeny which accommodate the well-established existence of admixed strains in this species (Liti *et al*., 2009; Peter *et al*., 2018). Despite using nearly twice as many strains, our phylogenetic tree of *S. cerevisiae* strains shows a similar topology as Peter *et al*. (2018), who identified a complex population structure including more than 26 distinct lineages plus many mosaic strains derived from admixture between these lineages. Strains in our integrated dataset that are not present in Peter *et al*. (2018) – such as those from Duan *et al*. (2018) and Barbosa *et al*. (2016) – cluster with known lineages previously characterized by Peter *et al*. (2018). For instance, “activated dry yeast” strains from Duan *et al*. (2018) cluster in the “mixed origin” lineage from Peter *et al*. (2018) (Figure 2).

**Figure 2:**
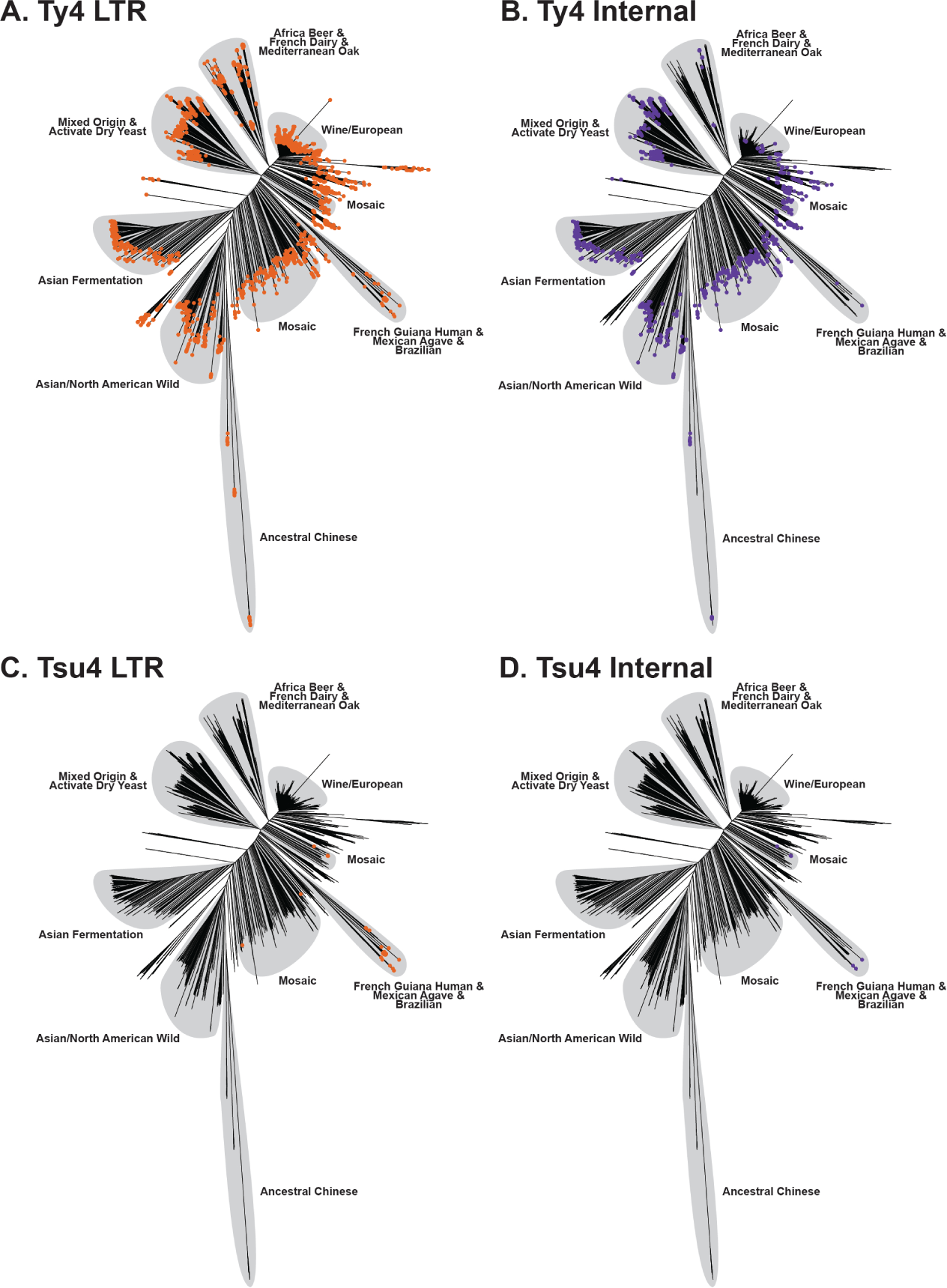
Host phylogeny of *S. cerevisiae* annotated with the presence or absence of Ty4/Tsu4 subfamilies. (A-D) NJ phylogenetic tree reconstructed using 2,787,577 genome-wide SNPs from 2,404 *S. cerevisiae* strains (see Materials and Methods). Major lineages are annotated with light gray shading based on previously-reported population structure (Peter *et al*., 2018; Duan *et al*., 2018). Colored dots indicate the presence of Ty4 (panels A and B) and Tsu4 (panels C and D) sequences for each *S. cerevisiae* strain. The presence of Ty4/Tsu4 subfamilies in a strain was inferred when copy number estimates were *>*1 for LTRs (annotated with orange dots in panel A and C) and *>*0.5 for internal regions (annotated with purple dots in panel B and D).

We then visualized the presence/absence of LTR and internal regions for Ty4/Tsu4 subfamilies on the phylogeny inferred for *S. cerevisiae* from short-read WGS data. This analysis revealed that Ty4 LTR and internal sequences are present in all *S. cerevisiae* lineages (Figure 2A,B). In contrast, Tsu4 LTR sequences are restricted to *∼*2% of strains surveyed (49/2,404) all of which are found in Central/South America (specifically French Guiana, Mexico, Brazil, and the French West Indies) (Figure 2C,D). Tsu4 sequences are completely absent from most other *S. cerevisiae* lineages, including the most ancestral Chinese lineages (Wang *et al*., 2012; Duan *et al*., 2018). We identified six *S. cerevisiae* strains that contain evidence of Tsu4 internal regions (245, AFQ and CDM from Mosaic lineage 2; CQS from the French Guiana lineage; UFMG-CM-Y641 and UFMG-CM-Y642 from the Brazil 3 lineage) (Marsit *et al*., 2015; Peter *et al*., 2018; Barbosa *et al*., 2016). Three of these *S. cerevisiae* strains (AFQ, CDM and CQS) also contain internal regions for Ty4.

Next, we analyzed Ty4/Tsu4 subfamily content in WGAs for a global sample of 183 *S. cerevisiae* strains (Table S1 and Figure S1), which confirmed results based on short-read WGS data. Ty4 subfamily sequences were found in all *S. cerevisiae* WGAs analyzed, while Tsu4 subfamily sequences were absent from the majority of *S. cerevisiae* WGAs. CQS is the only strain assembled using long-read data for which we identify FLEs for Tsu4 (n=9), confirming previous observations (O’Donnell *et al*., 2023). We also identified one full-length (245 and AFQ) or truncated (CDM) Tsu4 copy in short-read WGAs (Table S1) for three *S. cerevisiae* strains that we identified previously as containing Tsu4 internal regions in the short-read WGS scan. No publicly-available WGAs are available for the two Brazil 3 strains (UFMG-CM-Y641 and UFMG-CM-Y642, (Barbosa *et al*., 2016)) with evidence of Tsu4 internal regions based on WGS data. All three *S. cerevisiae* strains with Tsu4 FLEs in WGAs are geographically restricted to Central/South America. Importantly, we note that FLEs for both the Ty4 and Tsu4 subfamilies were identified in the long-read WGA for strain CQS.

The prevalence of the Ty4 subfamily in most *S. cerevisiae* lineages – including ancestral Chinese lineages (Wang *et al*., 2012; Duan *et al*., 2018) – indicates that the Ty4 subfamily was present in the MRCA of this species. However, despite being broadly active at the species level, the absence of Ty4 internal regions and FLEs in many strains indicates this subfamily has undergone many local extinction events (see also (Bleykasten-Grosshans *et al*., 2021)). In contrast, the absence of Tsu4 in most lineages (including ancestral Chinese lineages) strongly indicates that this subfamily was not present in the MRCA of *S. cerevisiae*. The small number of strains that do contain Tsu4 in *S. cerevisiae* do not form a single monophyletic group, which is consistent either with one HTT event followed by admixture among lineages, or multiple recent HTT events that have introduced Tsu4 into different lineages of *S. cerevisiae* in Central/South America. Finally, the observation of *S. cerevisiae* strains with FLEs for both Tsu4 and Ty4 subfamilies (i.e., CQS) demonstrates that FLEs from the Ty4 and Tsu4 subfamilies can co-exist in the same *Saccharomyces* strain.

### Multiple HTT events have transferred Tsu4 into *S. paradoxus* and *S. cerevisiae*

The short-read WGS strategy used above allowed us to establish Ty4 as the ancestral subfamily in *S. paradoxus* and *S. cerevisiae*, and to identify at least one Tsu4 HTT event in both species. However, this approach cannot resolve how many Tsu4 HTT events occurred in either species, nor can it identify the potential donor lineages for these HTT events. To investigate whether the presence of Tsu4 in *S. paradoxus* and *S. cerevisiae* can be explained by one or more Tsu4 HTT event, and to identify the most likely donor lineage(s) for these HTT events, we analyzed the molecular evolution of all Ty4 family FLEs identified in a integrated dataset of 210 high-quality WGAs for all recognized species in the genus *Saccharomyces*. The majority of WGAs in this dataset were generated using PacBio or Oxford Nanopore Technology (ONT) long reads, including new high-quality WGAs for *S. mikatae* strains IFO 1815 and NBRC 10994 that we generated using PacBio long reads (Table S2). Phylogenetic analysis of our new *S. mikatae* WGAs confirmed the taxonomic placement of IFO1815 in the Asia A clade (Peris *et al*., 2023) and revealed that NBRC 10994 should be placed in a new clade (Asia C) (Figure S2). In addition, we also included WGAs based on short-read data for three *S. cerevisiae* strains with Tsu4 internal sequences (245, AFQ and CDM; see above) and for two strains of *S. arboricola* (H-6 and ZP960) that represent the best available WGAs for this species.

In total, we identified 247 FLEs for the Ty4 subfamily and 124 FLEs for the Tsu4 subfamily in this integrated dataset (Table S2, Figure S1). No FLEs for either subfamily were identified *S. arboricola* and *S. kudriavzevii* using our current query sequences. The absence of Ty4 family FLEs in *S. arboricola* may simply reflect the lack of high-quality WGAs for this species. However, the absence of Ty4 family FLEs in *S. kudriavzevii* is likely an artifact of divergence between our current query sequences and Ty4 family FLEs in this species. In *S. kudriavzevii* strain (IFO1802) we observed a high copy number of truncated Tsu4 elements (n=8) that, upon further inspection, revealed five nearly full-length elements that were highly similar to one another, dispersed throughout the IFO1802 genome, and overlapped full-length *de novo* LTRharvest predictions (Ellinghaus *et al*., 2008). We concluded that these five *S. kudriavzevii* elements represented FLEs from a novel branch in the Ty4 family and included them in our phylogenetic analysis of FLEs.

We next created a multiple sequence alignment and reconstructed phylogenetic networks and trees based on internal coding regions of all 376 FLEs in our integrated dataset (Figure 3A, B). We excluded LTR and untranslated sequences from this analysis, which exhibited poor alignment due to higher divergence in noncoding regions. This analysis identified 14 well-supported clades of FLEs each found in a single species, plus two branches with singleton FLEs from the Hawaiian *S. paradoxus* strain UWOPS91-917.1 and Asia C *S. mikatae* strain NBRC 10994, respectively. Two clades (Clades 11 and 12) with FLEs from either *S. mikatae* or *S. kudriavzevii* exhibit evidence of reticulation between the Ty4 and Tsu4 clades (Figure 3A), which we interpret as being caused by recombination between these subfamilies (see detailed analysis below). Exclusion of these clades eliminated the major signal for reticulation between the Ty4 and Tsu4 subfamilies in the phylogenetic network (Figure S3A) and increased bootstrap support for clades in *S. jurei* and *S. mikatae*, but did not alter the topological relationships of other clades in the ML tree (Figure S3B). The Ty4 subfamily is represented by only two clades with FLEs from only *S. cerevisiae* (Clade 13) or *S. paradoxus* (Clade 14), respectively. In contrast, the Tsu4 subfamily is represented by ten species-specific clades (1-10) with FLEs from all species except *S. kudriavzevii* (Figure 3A, S4, S5, S6). Two clades each of Tsu4 FLEs from *S. paradoxus* (Clades 1 and 3) and *S. cerevisiae* (Clades 2 and 4) together with the singleton FLE from *S. paradoxus* UWOPS91-917.1 form a monophyletic group. *S. eubayanus* is represented two Tsu4 FLEs clades from the Holarctic (Clade 5) and Patagonian (Clade 6) lineages, respectively. Tsu4 FLEs from *S. uvarum* form two clades (Clades 7 and 8) that are both found in a single strain from the Holarctic lineage. Tsu4 FLEs from European *S. jurei* (Clade 9) cluster with Tsu4 FLEs from the *S. mikatae* Asia A lineage (Clade 10) and the singleton branch from the *S. mikatae* Asia C lineage.

**Figure 3:**
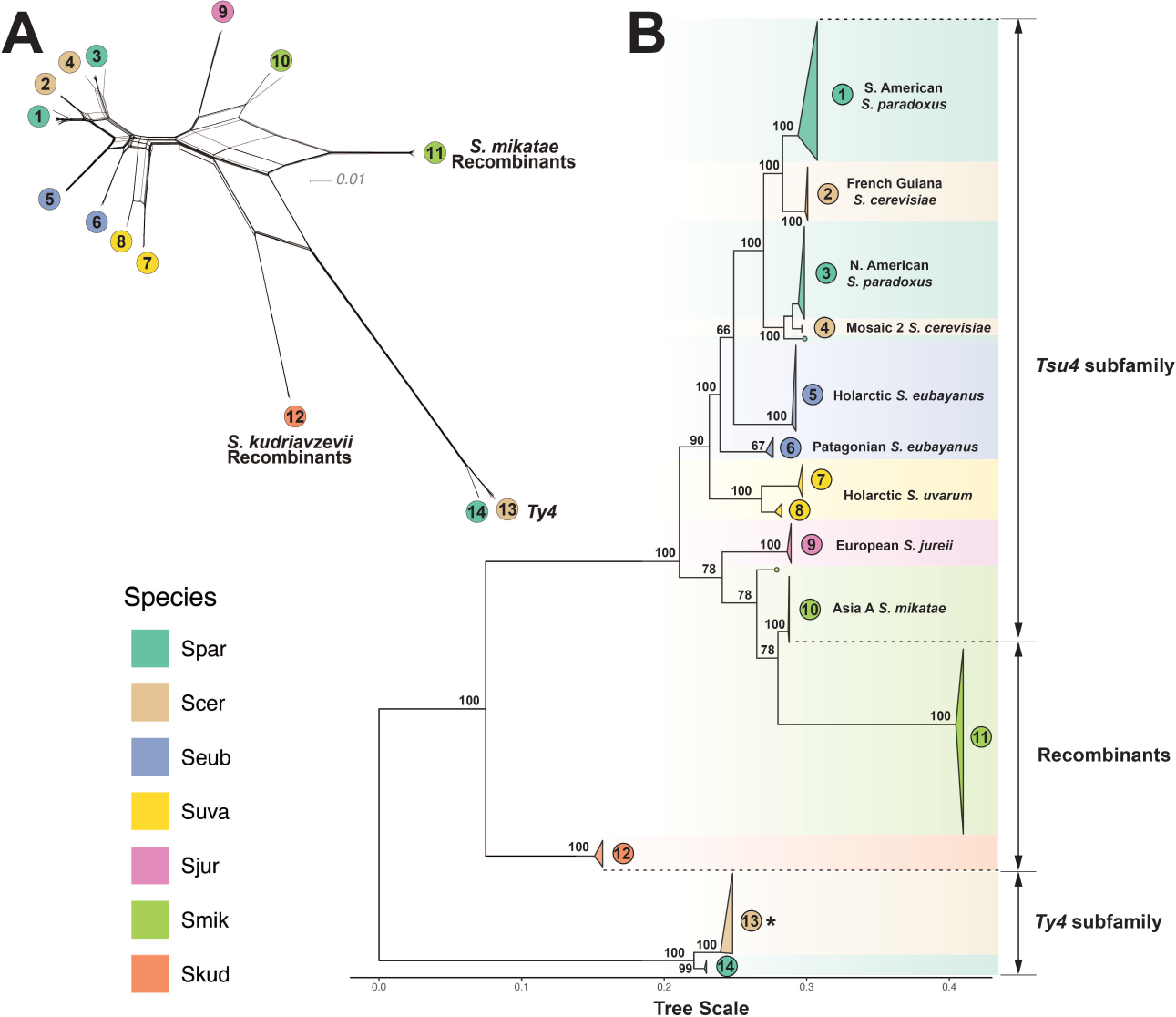
Phylogenetic network and tree of FLEs from the Ty4 family in *Saccharomyces*. (A) Phylogenetic network for internal coding regions of Ty4/Tsu4 FLEs based on the NeighborNet algorithm. To simplify visualization, this network only includes Ty4 subfamily FLEs from WGAs reported in Yue *et al*. (2017). Lineages in the network are labeled according to monophyletic groups identified in panel B. (B) Midpoint rooted ML phylogeny of internal coding regions from Ty4/Tsu4 FLEs. Bootstrap support based on 100 replicates is shown for major nodes. The scale bar for branch lengths is in units of substitutions per site. All monophyletic groups are collapsed as triangles. Two singleton Tsu4 elements (f267 from Hawaiian *S. paradoxus* strain UWOPS91-917.1 and f256 from *S. mikatae* strain NBRC 10994) are denoted as dots at tips. Triangles, tip dots, and ranges are colored for each species. Vertical heights of triangles are proportional to the number of taxa. Horizontal widths of triangles are equal to the maximum branch length within the clade. Note that the monophyletic clade for the Ty4 subfamily from *S. cerevisiae* (annotated with an asterisk) is re-scaled to 5% of the real sample size both horizontally and vertically, due to the large number of Ty4 sequences (n=244) in *S. cerevisiae* genomes.

Previous analysis of Tsu4 HTT events in *S. paradoxus* and *S. cerevisiae* using a smaller dataset of FLEs (Bergman, 2018) suggested one primary HTT occurred in the ancestor of American *S. paradoxus* lineages followed by one secondary HTT from *S. paradoxus* into *S. cerevisiae*. This hypothesis predicts that Tsu4 FLEs from *S. paradoxus* and *S. cerevisiae* should form a single clade with *S. cerevisiae* FLEs forming a single sub-clade somewhere within the broader diversity of *S. paradoxus* FLEs. Two key features of the Tsu4 FLE phylogeny in our expanded dataset are inconsistent with this hypothesis (Figure 3). First, we observe two distinct clades of FLEs for both *S. paradoxus* and *S. cerevisiae*, each of whose closest sampled relatives are from a different species. Namely, *S. cerevisiae* Clade 2 (from the French Guiana strain CQS) clusters with S. American *S. paradoxus* Clade 1, while *S. cerevisiae* Clade 4 (from Mosaic 2 strains 245 and AFQ) clusters with the N. American *S. paradoxus* Clade 3. Second, the observed topology of Tsu4 FLEs in *S. paradoxus* does not strictly follow the host phylogeny for the SpB sub-lineage strain UFRJ50816. Specifically, Tsu4 FLEs from the S. American *S. paradoxus* strain UFRJ50816 form their own divergent Clade 1 instead of grouping as expected with Tsu4 FLEs from other SpB strains from N. America (MSH-604 and YPS138) in Clade 3 (Figure S4).

We interpret the phylogeny of Tsu4 FLEs in *S. paradoxus* and *S. cerevisiae* to be consistent with at least two Tsu4 HTT events into each species. In *S. paradoxus*, we infer one ancestral Tsu4 HTT event creating Clade 3 FLEs that are present in all SpB, SpC, SpC* and SpD strains (except UFRJ50816), and one recent HTT event creating Clade 1 FLEs present in the S. American sub-lineage containing UFRJ50816 (where Clade 3 FLEs had already gone extinct). Likewise, in *S. cerevisiae*, we infer one recent HTT into the French Guiana lineage creating Clade 2 FLEs, and another recent HTT into the Mosaic 2 lineage creating Clade 4 FLEs. These data suggest that when conditions are favorable for HTT events in *Saccharomyces*, they can occur more frequently than the most parsimonious interpretation based on presence/absence data would imply.

### Tsu4 in *S. paradoxus*, *S. cerevisiae*, and Holarctic *S. eubayanus* were transferred from an unknown donor

The collective monophyly of the Tsu4 clades found in *S. paradoxus* and *S. cerevisiae* suggests the Tsu4 HTT events in these species ultimately arose from a similar donor lineage, with distinct clades being formed at different times or in different geographic regions. Using indirect data from the hybrid species *S. pastorianus* that contains subgenomes from *S. eubayanus* and *S. cerevisiae* (Libkind *et al*., 2011; Baker *et al*., 2015; Okuno *et al*., 2016), Bergman (2018) concluded that the Holarctic lineage of *S. eubayanus* contains the most closely related Tsu4 sequences to those in *S. paradoxus* and *S. cerevisiae*. Our current analysis provides direct evidence for this conclusion, with Clade 5 FLEs from CDFM21L.1 in the the Holarctic *S. eubayanus* lineage clustering most closely with the common ancestor of all Tsu4 FLEs in *S. paradoxus* and *S. cerevisiae* (Figure 3). Taken at face value, this result implies that Holarctic *S. eubayanus* represents the most likely donor lineage for the multiple HTT events observed in *S. paradoxus* and *S. cerevisiae*.

However, several features of the Tsu4 FLE phylogeny suggest that the Holarctic *S. eubayanus* lineage is not the direct donor for the HTT events in *S. paradoxus* and *S. cerevisiae* (Figure 3). First, Tsu4 clades in *S. paradoxus* and *S. cerevisiae* are not nested within the diversity of FLEs from Holarctic *S. eubayanus*, but rather form a sister group separated by substantial divergence. Second, bootstrap support for the clustering of Holarctic *S. eubayanus* FLEs with the ancestor of FLEs from *S. cerevisiae* and *S. paradoxus* is relatively weak (*>*66%). The alternative clustering of FLEs from the Holarctic and Patagonia-B *S. eubayanus* lineages together would suggest that the donor into *S. paradoxus* and *S. cerevisiae* is from a currentlyunsampled lineage of *S. eubayanus*, or a species closely related to *S. eubayanus*. Third, Tsu4 FLEs from Holarctic *S. eubayanus* are not nested with in the diversity of *S. eubayanus* Patagonian FLEs (Figure 3, S5), as is expected since Holarctic *S. eubayanus* is known to be a sub-lineage of the Patagonia-B lineage (Figure S7) (Peris *et al*., 2016). Discordance between the *S. eubayanus* Tsu4 and host strain phylogenies suggests the possibility of a previously-undetected Tsu4 HTT event into Holarctic *S. eubayanus*, which could lead to the false conclusion that the Holarctic *S. eubayanus* lineage is the most likely donor for HTT events into *S. paradoxus* and *S. cerevisiae*.

To test for a previously-undetected Tsu4 HTT event in the Holarctic *S. eubayanus* lineage, we developed a novel approach to study Tsu4 sequence evolution using strain-specific consensus sequences inferred from short-read WGS data. Importantly, this approach bypasses the limited number of WGAs available in *S. eubayanus* and other potential donor species and allows us to generalize results across larger samples of host strains and lineages. The premise behind this approach is based on the observation that Tsu4 FLEs typically cluster first within the same strain before clustering with FLEs from other strains (Figure 3). Thus, strainspecific Tsu4 consensus sequences should be a reasonable proxy for the common ancestor of elements within a strain, and can themselves be used for evolutionary inference across strains and species.

Using the short-read based WGS approach as above for *S. paradoxus* and *S. cerevisiae*, we first estimated Ty4/Tsu4 LTR and internal copy numbers in the context of host phylogenies for *S. eubayanus* (Figure S7) and *S. uvarum* (Figure S8), respectively. These results reveal that Tsu4 was present and Ty4 was absent in the ancestors of both *S. eubayanus* and *S. uvarum*, that Tsu4 is broadly active in both species, and that the Holarctic *S. eubayanus* lineage has the highest LTR copy number of Tsu4 in either species. We then computed consensus sequences for Tsu4 internal regions in all *S. paradoxus*, *S. cerevisiae*, *S. eubayanus* and *S. uvarum* strains with an estimated copy number of *>*0.75 and generated a ML tree of strain-specific sequences across the combined dataset of four species (Figure 4).

**Figure 4:**
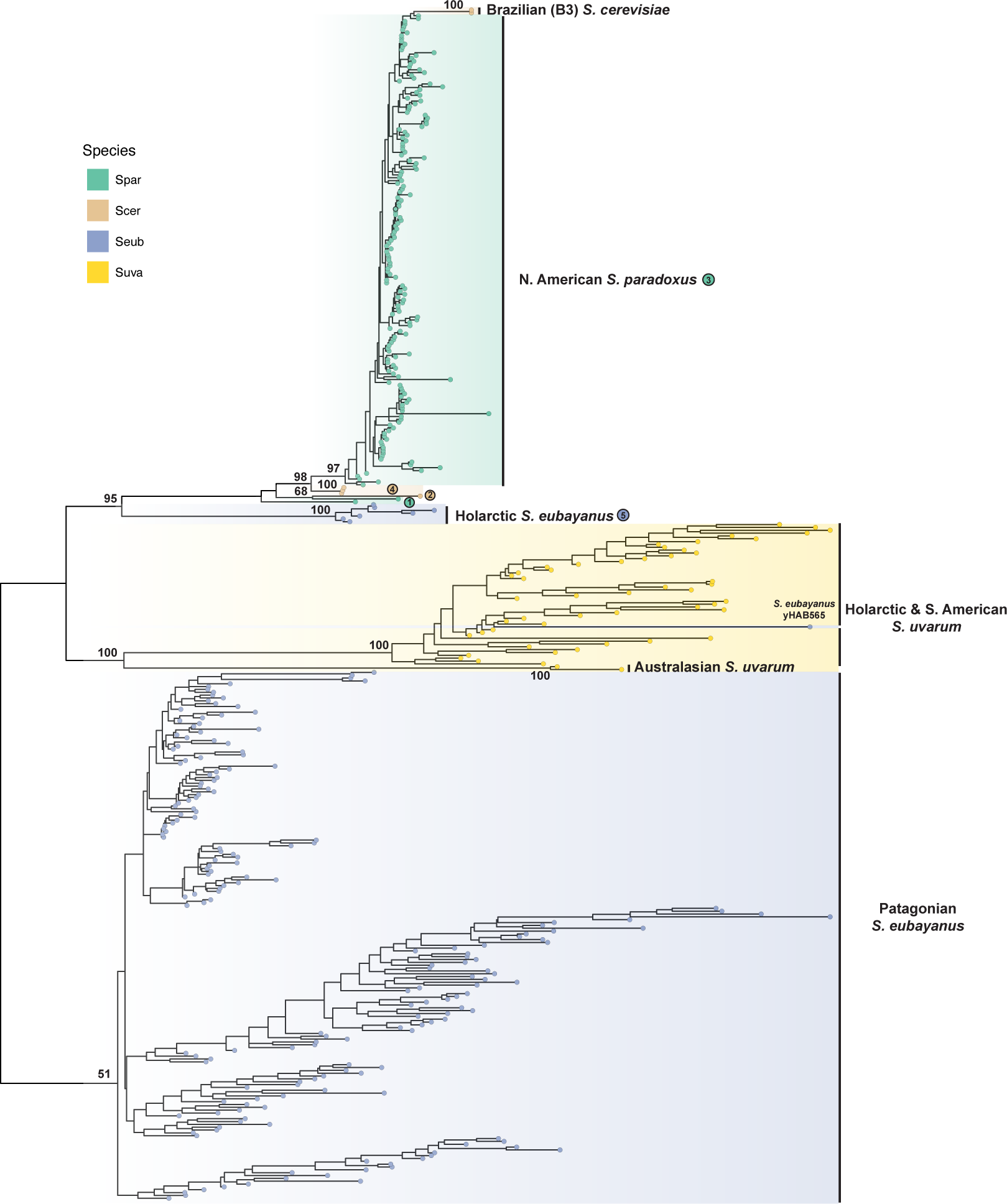
ML phylogeny of strain-specific consensus sequences for Tsu4 internal regions in all *S. paradoxus*, *S. cerevisiae*, *S. eubayanus* and *S. uvarum*. Shown is the phylogeny reconstructed with 1,817 distinct alignment sites from 419 strain-specific consensus sequences. The consensus sequence is computed for each *S. paradoxus*, *S. cerevisiae*, *S. eubayanus*, and *S. uvarum* strain that has *>*0.75 depth and *>*0.9 breadth in its Tsu4 internal region. Tip points are colored by species. The phylogeny is midpoint rooted. Bootstrap supporting values are annotated for key nodes. Key clades are annotated with host lineage and/or clade numbers from the Tsu4 FLE phylogeny.

Key groupings in the strain-specific Tsu4 consensus tree (Figure 4) agreed with the phylogeny of Tsu4 FLEs based on a smaller number of strains above (Figure 3), cross-validating both approaches. Notably, Tsu4 consensus sequences for all N. American *S. paradoxus* strains (together with the two *S. cerevisiae* Brazil 3 strains, UFMG-CM-Y641 and UFMG-CM-Y642) form a large monophyletic group (corresponding to Clade 3 in the FLE tree) that clusters most closely with consensus sequences from Mosaic 2 *S. cerevisiae* strains (corresponding to Clade 4). Likewise, the consensus sequence for the S. American *S. paradoxus* strain UFRJ50916 (corresponding to Clade 1) clusters with the S. American *S. cerevisiae* strain CQS (corresponding to Clade 2). All seven Holarctic *S. eubayanus* strains form a monophyletic group (corresponding to Clade 5) that clusters with *S. paradoxus* and *S. cerevisiae* consensus sequences and is distinct from Tsu4 sequences found in all other *S. eubayanus* or *S. uvarum* strains.

Phylogenetic analysis of strain-specific consensus sequences also revealed two other Tsu4 HTT events that could not be detected in the FLE phylogeny because of limited samples of WGAs. The first involves the two *S. cerevisiae* Brazil 3 strains (UFMG-CM-Y641 and UFMG-CM-Y642). Based on the location of Brazil 3 samples in the consensus sequence tree (Figure 4) and the pattern of variation in the consensus sequences of all six *S. cerevisiae* strains that have Tsu4 (Figure S9), we conclude that Tsu4 sequences in the *S. cerevisiae* Brazil 3 lineage arose from a HTT event that is distinct from those detected using FLEs in the Mosaic 2 or French Guiana lineages. The second peviously-undetected Tsu4 HTT event involves the *S. eubayanus* Patagonia B strain yHAB565, whose Tsu4 consensus sequence is placed within the *S. uvarum* cluster. *S. eubayanus* yHAB565 is placed correctly in the Patagonia B lineage in our host strain tree (Figure S7), which rules out the possibility of sample mixups during sequencing or bioinformatic analysis and supports a recent HTT Tsu4 event from *S. uvarum* into *S. eubayanus*.

Taken together, our results suggest that the similarity between the Tsu4 clades in Holarctic *S. eubayanus*, *S. paradoxus*, and *S. cerevisiae* arose from parallel HTT events donated by a common but as-yet-unidentified *Saccharomyces* lineage. The substantial divergence between non-Holarctic *S. eubayanus* or *S. uvarum* and the clade comprised of Holarctic *S. eubayanus*, *S. paradoxus* and *S. cerevisiae* suggests that this unknown donor is either a currently-unsequenced lineage of *S. eubayanus* or *S. uvarum* (e.g., West China *S. eubayanus* (Bing *et al*., 2014)) or potentially an undiscovered species related to *S. eubayanus* and *S. uvarum*. Additionally, we show that our novel strain-specific consensus sequence approach complements analysis of FLEs from WGAs and can reveal previously undetected cases of Tsu4 HTT in the abundant WGS datasets that are available in multiple yeast species.

### Tsu4 HTT fuels the evolution of recombinant clades in *S. mikatae* and *S. kudriavzevii*

Our phylogenetic network analysis of FLEs from the Ty4 family above revealed evidence of reticulation in *S. mikatae* Clade 11 and *S. kudriavzevii* Clade 12 (Figure 3A) that could be caused by recombination between the Ty4 and Tsu4 subfamilies (Huson and Bryant, 2006). In addition, the coexistence of FLEs for both Ty4 and Tsu4 in *S. cerevisiae* strain CQS indicates that conditions for recombination between Ty4 and Tsu4 subfamilies can occur in nature. To provide further evidence for recombination between the Ty4 and Tsu4 subfamilies, we first selected representative FLEs for “pure” Tsu4 (f32 from the *S. uvarum* Clade 8) and “pure” Ty4 (f49 from the *S. cerevisiae* Clade 13) from outgroup species not involved in the putative recombination events. We then plotted a sliding window of pairwise sequence divergence between these representative pure Ty4 and Tsu4 FLEs and a putatively “pure” *S. mikatae* Clade 10 FLE (f286 from IFO 1815) or a putatively “recombinant” *S. mikatae* Clade 11 FLE (f256 from IFO 1815) (Figure S10). This analysis revealed that that the 5’ internal region – including the complete *gag* gene and the first *∼*500bp of *pol* – shows lower levels of divergence between *S. uvarum* Tsu4 and pure *S. mikatae* Clade 10 (Figure S10A) than recombinant *S. mikatae* Clade 11 (Figure S10B). Conversely, the same 5’ internal region shows higher levels of divergence between *S. cerevisiae* Ty4 and pure *S. mikatae* Clade 10 (Figure S10D) than recombinant *S. mikatae* Clade 11 (Figure S10E). These data indicate that the 5’ internal segment in Clade 11 is derived from the Ty4 subfamily, while the rest of the Clade 11 internal region is derived from the Tsu4 subfamily.

We then partitioned the multiple sequence alignment of Ty4 family FLE internal regions into 5’ and 3’ segments, and reconstructed ML phylogenies for both partitions from representative clades (Figure 5). A striking discordance can be observed in phylogenies reconstructed from 5’ and 3’ internal regions of Clade 11 sequences. In the 5’ partition, pure Clade 10 FLEs cluster with Tsu4 FLEs from *S. jurei* and *S. uvarum*, while recombinant Clade 11 FLEs cluster with strong support as a sister group to from *S. paradoxus*/*S. cerevisiae* in the Ty4 subfamily (Figure 5A). In contrast, in the tree reconstructed from the 3’ partition *S. mikatae* Clades 10 and 11 form a single monophyletic group that is closely related to Tsu4 sequences from *S. jurei* (Figure 5B).

**Figure 5:**
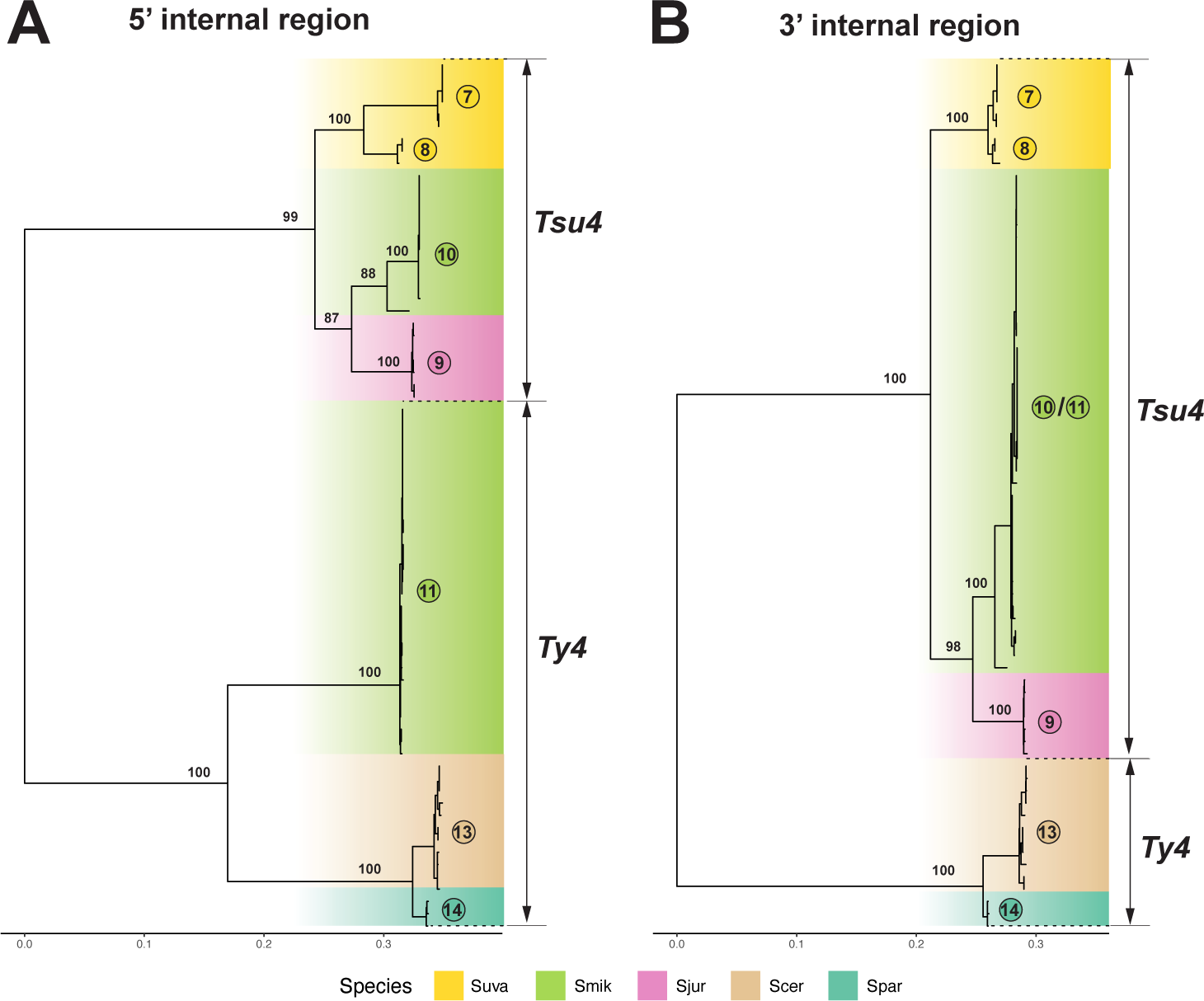
ML phylogeny for 5’ and 3’ internal regions from Tsu4 FLEs in *S. mikatae*, *S. jurei* , and *S. uvarum*, plus representatives of Ty4 elements. Panel (A) shows the ML phylogeny for 5’ internal region containing 459 distinct alignment sites; Panel (B) shows 3’ internal region containing 455 distinct alignment sites. Internal coding regions from 71 Ty4 and Tsu4 FLEs are included in both panels. Both trees are midpoint rooted, and visualized in the same tree scale which is shown in units of substitutions per site. Bootstrap supporting values are annotated for key nodes.

Based on these results and presence of Ty4 solo LTRs in WGAs from *S. mikatae* and *S. jurei* (Table S1), we propose the following scenario for the evolution of *S. mikatae* Clades 10 and 11. A divergent Ty4 subfamily was previously active in an ancestor of *S. mikatae* and *S. jurei* , which has subsequently gone extinct in both species but left Ty4 internal sequences that were retained in the *S. mikatae* genome for some period of time. A HTT event introduced the Tsu4 subfamily prior to the speciation of *S. mikatae* and *S. jurei* , which evolved into Clade 10 in *S. mikatae* and Clade 9 in *S. jurei* . This Tsu4 HTT event explains the discordance between the Ty4 family and host species phylogenies previously reported for *S. mikatae* Bergman (2018). The donor for this Tsu4 HTT into the ancestor of *S. mikatae* and *S. jurei* is unknown but related to the ancestor to all extant Tsu4 FLEs in *S. eubayanus*, *S. uvarum*, *S. paradoxus* and *S. cerevisiae*. Recombination of the 5’ internal region from the now-extinct Ty4 subfamily in *S. mikatae*onto a Clade 10-like pure Tsu4 FLE created the “recombinant” Tsu4 Clade 11. This model explains the lack of Ty4 FLEs in *S. mikatae* and *S. jurei* (Table S1), the coexistence of two highly divergent clades in *S. mikatae* genomes (Figure 3B), the long branch leading to Clade 11 in the FLE phylogeny (Figure 3B), and reticulation between Ty4 and Tsu4 subfamilies for Clade 11 in the phylogenetic network (Figure 3A).

We applied similar approaches to investigate whether reticulation in the phylogenetic network observed for *S. kudriavzevii* Clade 12 FLEs (Figure 3A) also is caused by recombination between Ty4 and Tsu4 subfamilies. Sliding window analysis of a representative Clade 12 FLE versus pure Tsu4 and Ty4 from outgroup species revealed an *∼*2kb segment starting at the beginning of Pol that shows very high similarity to Tsu4 (Figure S10C). Phylogenetic analysis of partitions corresponding to the “left,” “middle,” and “right” segments of FLE internal regions for representative clades reveals that the middle internal segment of Clade 12 is derived from the Tsu4 subfamily, while the left and right segments are divergent representatives of the Ty4 subfamily. Based on these results and presence of Ty4 LTRs in WGAs from *S. kudriavzevii* (Table S1), we propose that *S. kudriavzevii* ancestrally contained a divergent Ty4 subfamily which acquired a middle segment from a horizontally-transferred Tsu4 by recombination. The Tsu4 clade that was horizontally transferred into *S. kudriavzevii* and the original pure *S. kudriavzevii* Ty4 clade have both subsequently gone extinct, leaving the recombinant Clade 12 as the only extant representative of the Ty4 family currently identified in *S. kudriavzevii* . Based on clustering of the middle internal region (Figure 6B), the donor lineage for the Tsu4 HTT into *S. kudriavzevii* is related to the donor for Tsu4 HTT in *S. mikatae* and *S. jurei* . Together, the recombinant clades in *S. mikatae* and *S. kudriavzevii* support the conclusion that co-existence of Ty4 and Tsu4 subfamily sequences in the same genome mediated by HTT provides substrate for recombination to generate new retrotransposon clades in *Saccharomyces*.

**Figure 6:**
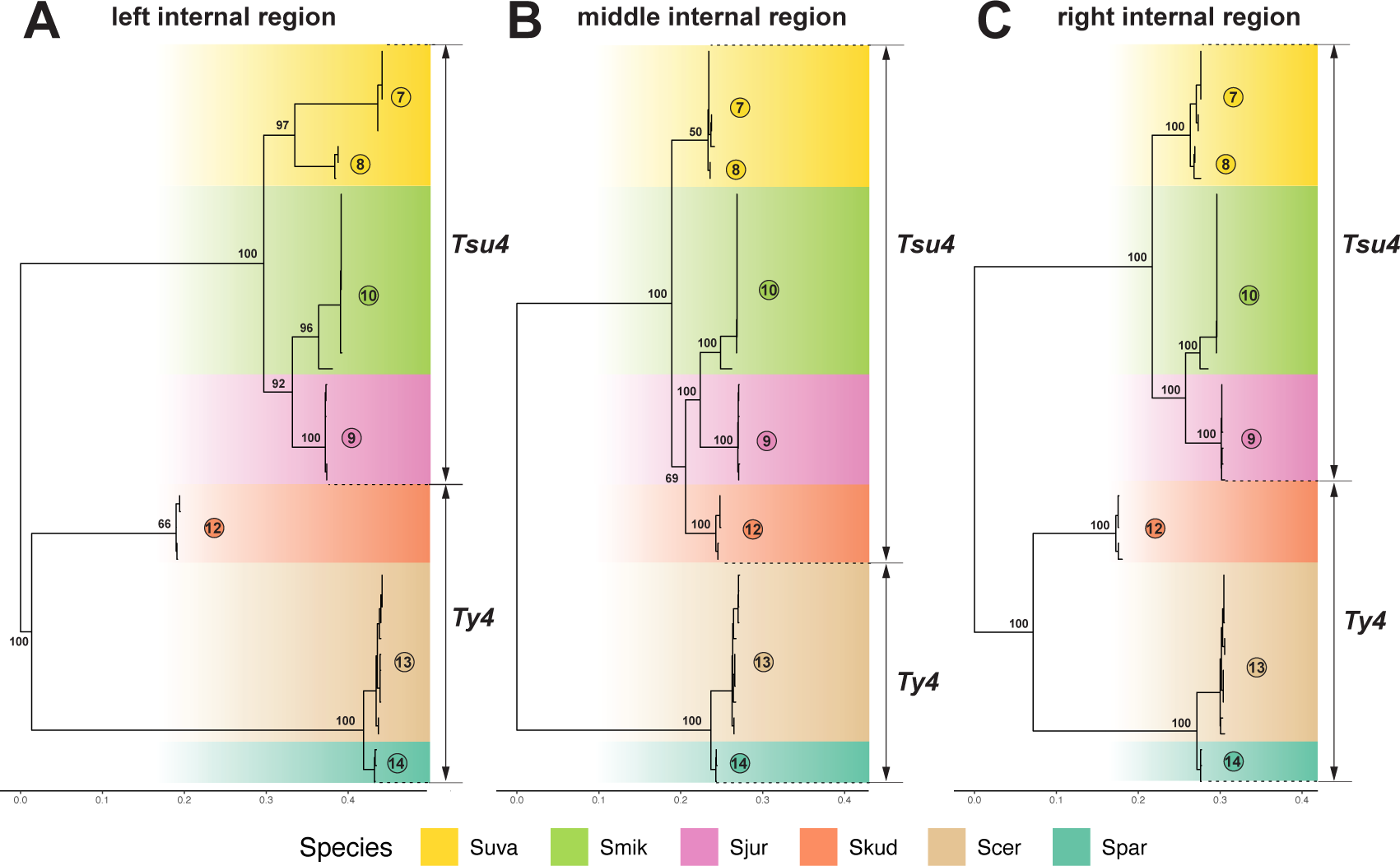
ML phylogeny for partitioned internal regions from recombinants in *S. kudriavzevii* , pure Tsu4 FLEs in *S. mikatae*, *S. jurei* , and *S. uvarum*, plus representatives of Ty4 elements. Panel (A) shows the ML phylogeny for left-side internal region based on 357 distinct alignment sites; Panel (B) for middle internal region based on 367 distinct alignment sites; Panel (C) for right-side internal region based on 423 distinct alignment sites. In all three panels, internal regions from 47 Ty4 and Tsu4 FLEs are included. All trees are midpoint rooted, and visualized in the same tree scale which is shown in units of substitutions per site. Bootstrap supporting values are annotated for key nodes.

## Conclusions

Here we address open questions concerning the impact of HTT on the evolution of the Ty4 family in *Saccharomyces* by integrating large-scale short-read WGS data and high-quality long-read WGAs from multiple *Saccharomyces* species. We show that the previously detected Tsu4 HTT event in *S. paradoxus* (Bergman, 2018) occurred in the ancestor of all American lineages and report new evidence for a second recent Tsu4 HTT in the South American lineage of *S. paradoxus*. We also show that the previously reported presence of Tsu4 in *S. cerevisiae* (Bergman, 2018; O’Donnell *et al*., 2023) is explained by at least three independent recent HTT events into *S. cerevisiae* in Central/South America, at least one of which (into the French Guiana lineage) is also associated with HTT of another retrotransposon family (Ty1) and introgression of host genes from *S. paradoxus* (Peter *et al*., 2018; Bleykasten-Grosshans *et al*., 2021). We confirm that the Holarctic lineage of *S. eubayanus* contains Tsu4 elements that are most closely related to those in *S. paradoxus* and *S. cerevisiae* (Bergman, 2018) but conclude that this similarity is caused by an independent Tsu4 HTT into the Holarctic *S. eubayanus* lineage from an unidentified donor. Recurrent HTT of Tsu4 into different host species and lineages provides a mechanism to explain why this subfamily is more clade-rich than the Ty4 subfamily in the Ty4 family phylogeny.

Additionally, we investigate the putative Tsu4 HTT event reported for *S. mikatae* (Bergman, 2018) by generating new PacBio WGAs for two strains in this species (IFO 1815 and NBRC 10994), which revealed the presence of two active Ty4 family clades in *S. mikatae*. The first is a pure Tsu4 clade that clusters with FLEs from *S. jurei* , providing evidence for an ancestral Tsu4 HTT event prior to the divergence of *S. mikatae* and *S. jurei* that explains the discordance between Ty4 family and host species phylogenies for *S. mikatae* (Bergman, 2018). We also find a second recombinant clade in *S. mikatae* that shares similarity to the Ty4 and Tsu4 subfamilies in different parts of the internal region. The recombinant Clade 11 implies the co-existence of internal sequences for both subfamilies in the *S. mikatae* genome at some point in history. Likewise, we identify a novel clade in *S. kudriavzevii* that similarly exhibits recombination between the Ty4 and Tsu4 subfamilies, and explains the divergent position of *S. kudriavzevii* FLEs in the Ty4 family phylogeny (Bergman, 2018).

The discovery of novel Ty4 family clades in *S. mikatae* and *S. kudriavzevii* that were generated by recombination between resident (ancestral) and horizontally transferred (derived) retrotransposon subfamilies can be used to generalize prior results reported for the Ty1/Ty2 superfamily in *S. cerevisiae* (Jordan and McDonald, 1998; Czaja *et al*., 2020; Bleykasten-Grosshans *et al*., 2021). The canonical Ty1 subfamily found in *S. cerevisiae* evolved from an ancestral Ty1’ subfamily by independently acquiring segments from horizontally-transferred European *S. paradoxus* Ty1 and Ty2 elements by recombination (Jordan and McDonald, 1998; Czaja *et al*., 2020; Bleykasten-Grosshans *et al*., 2021). Similarly, the Ty101 subfamily evolved in *S. cerevisiae* through recombination between the ancestral Ty1’ subfamily and a horizontal transferred South American *S. paradoxus* Ty1 element (Bleykasten-Grosshans *et al*., 2021). Together these results suggest that recombination among divergent subfamilies that co-occur in the same species because of HTT may be a common mechanisms for the evolution of new of new retrotransposon lineages in *Saccharomyces*. In addition, Ty1c and Ty101 in *S. cerevisiae* and the recombinant Clade 11 in *S. mikatae* both retain a complete *gag* gene from the horizontally-transferred element. A truncated product from Ty1c *gag* has been shown to encode a copy-number dependent repressor of Ty1c transposition in *S. cerevisiae* and *S. paradoxus* (Saha *et al*., 2015; Cottee *et al*., 2021). Thus, determining the functional significance of recombinant elements with respect to fitness or transposition control mechanisms may be fruitful areas for future research.

Finally, the observation of multiple independent Tsu4 HTT events in both *S. paradoxus* and *S. cerevisiae* reinforces prior observations of parallel HTT events involving the Ty1 family in different lineages of *S. cerevisiae* (Czaja *et al*., 2020; Bleykasten-Grosshans *et al*., 2021). Together, these results suggest that parallel HTT events in different parts of a species range may be a common occurrence in *Saccharomyces*. If so, the number of HTT events in *Saccharomyces* species cannot be reliably inferred from simple presence/absence data within species, and reconstructing the complex history of HTT events will require high-resolution phylogenetic data from large samples of FLEs or strain-specific consensus sequences. Similarly, the observation of parallel HTT events on short timescales for multiple yeast TE families suggests that large-scale surveys that detect HTT among distantly related taxa may underestimate the frequency of HTT in eukaryotic genome evolution (Peccoud *et al*., 2017, 2018).

## Materials and Methods

### DNA preparation, PacBio sequencing and genome assembly of *S. mikatae* strains IFO 1815 and NBRC 10994

To prepare DNA for PacBio sequencing, single colonies of strain IFO 1815 (NCYC 2888) and NBRC 10994 was inoculated in 7 ml yeast extract-peptone-dextrose (YPD) liquid broth and cultured for *∼*24 hours at 30*^◦^*C. DNA was isolated using the Wizard genomic DNA purification kit (Promega), and a PacBio library was prepared using the SMRTbell Express template prep kit (Pacific Biosciences), following the *>*15-kb size-selection protocol that includes Covaris g-TUBE shearing. PacBio sequencing was performed using the Sequel II instrument (sequencing kit v2.1). Whole genome assemblies were generated by performing Flye (v2.9) (Kolmogorov *et al*., 2019) with parameters “–pacbio-raw -g 12m.” Raw PacBio reads and genome assemblies of both *S. mikatae* strains have been submitted to NCBI under BioProject PRJNA934353.

### Genome sequence datasets

We compiled public short-read paired-end WGS datasets of multiple *Saccha-romyces* species to generate intraspecific phylogenies and survey Ty4/Tsu4 sub-family content within species. WGS datasets for each *Saccharomyces* species were cleaned and normalized using the following steps: (i) raw reads with identical BioSample accession, strain name and sequencing strategy (i.e., sequencing instrument and layout) were merged according to the metatable provided by NCBI SRA; (ii) for the NCBI BioSample accessions that have multiple records after merging, only the record with the highest sequencing depth was retained; and (iii) all samples with sequencing depth *<*10*×* were removed. After these quality control processes, the integrated WGS dataset used in this study includes 2,404 *S. cerevisiae* strains (Skelly *et al*., 2013; Bergstrom *et al*., 2014; Almeida *et al*., 2015; Marsit *et al*., 2015; Song *et al*., 2015; Strope *et al*., 2015; Barbosa *et al*., 2016; Drozdova *et al*., 2016; Gallone *et al*., 2016; Gayevskiy *et al*., 2016; Goncalves *et al*., 2016; Zhu *et al*., 2016; Coi *et al*., 2017; Istace *et al*., 2017; Kita *et al*., 2017; Maclean *et al*., 2017; Yue *et al*., 2017; Barbosa *et al*., 2018; Duan *et al*., 2018; Legras *et al*., 2018; Peter *et al*., 2018; Fay *et al*., 2019; Kang *et al*., 2019; Ramazzotti *et al*., 2019; Basile *et al*., 2021; Bigey *et al*., 2021; Han *et al*., 2021), 370 *S. paradoxus* strains (Bergstrom *et al*., 2014; Leducq *et al*., 2016; Xia *et al*., 2017; Eberlein *et al*., 2019; Koufopanou *et al*., 2020; He *et al*., 2022; Peris *et al*., 2023), 18 *S. mikatae* strains (Peris *et al*., 2023), 62 *S. uvarum* strains (Scannell *et al*., 2011; Almeida *et al*., 2014; Macias *et al*., 2021; Peris *et al*., 2023), and 292 *S. eubayanus* strains (Peris *et al*., 2016; Brouwers *et al*., 2019; Salazar *et al*., 2019; Langdon *et al*., 2020; Nespolo *et al*., 2020; Bergin *et al*., 2022; Mardones *et al*., 2022; Molinet *et al*., 2022; Peris *et al*., 2023).

We complied public high-quality WGAs of *Saccharomyces* species to identify Ty4/Tsu4 copies and extract FLEs for phylogenetic analysis. These WGAs were generated mostly with long-read sequencing data (PacBio or ONT), however we also included three WGAs generated with short-read sequencing data for *S. cerevisiae* that showed evidence of Tsu4 internal regions (strain 245 from (Marsit *et al*., 2015); strains AFQ and CDM from (Peter *et al*., 2018)) and two WGAs generated with short-read sequencing data for *S. arboricola* (strain H-6 from (Liti *et al*., 2013) and strain ZP960 from (Peris *et al*., 2023)) that represent the best available WGAs for this species. In total, we analyzed 183 *S. cerevisiae* WGAs (Marsit *et al*., 2015; Yue *et al*., 2017; Peter *et al*., 2018; O’Donnell *et al*., 2023), 12 *S. paradoxus* WGAs (Yue *et al*., 2017; Eberlein *et al*., 2019; Chen *et al*., 2022b), two *S. mikatae* WGAs (this study), two *S. jurei* WGAs (Naseeb *et al*., 2018), three *S. kudriavzevii* WGAs (Boonekamp *et al*., 2018; Salzberg *et al*., 2022), two *S. arboricola* WGAs (Liti *et al*., 2013; Peris *et al*., 2023), two *S. uvarum* WGAs (Chen *et al*., 2022a; Salzberg *et al*., 2022), and four *S. eubayanus* WGAs (Brickwedde *et al*., 2018; Brouwers *et al*., 2019; Mardones *et al*., 2022).

### Ty4/Tsu4 copy number estimates

Copy number of LTRs and internal regions for the Ty4 and Tsu4 subfamilies were estimated by the coverage module of McClintock 2 (Chen *et al*., 2023) using public short-read WGS datasets compiled above. For this analysis, we used McClintock revision 7aa5298 with parameters “--keep_intermediate minimal,coverage -m coverage”. Reference genomes used for WGS based copy number were as follows: *S. cerevisiae* laboratory strain S288c (UCSC version sacCer2); *S. paradoxus* European strain CBS432 (GCA_002079055.1) (Yue *et al*., 2017); *S. uvarum* European strain CBS 7001 (GCA_019953615.1) (Chen *et al*., 2022a); *S. eubayanus* Patagonia strain FM1318 (GCA_001298625.1) (Baker *et al*., 2015). The Ty query library used for this analysis is the same as in Czaja *et al*. (2020). The edge-trimming option in the McClintock coverage module was disabled by specifying “omit_edges” as “False” in the configuration file “config/coverage/coverage.py”. To reduce the influence of variable coverage and computing resources, samples with original fold-coverage greater than 100*×* were down-sampled to 100*×* using seqtk (v1.3) (https://github.com/lh3/seqtk).

### Phylogenetic analysis of host species

For each species, multi-sample variant calling was performed with BCFtools (v1.16, “bcftools mpileup -a ‘FORMAT/DP’ -Q 20 -q 20”; “bcftools call -f GQ,GP -mv – skip-variants indels”) (Li, 2011) using BAM files generated by McClintock 2 (Chen *et al*., 2023). Alignments with mapping quality less than 20, or bases with quality score less than 20, were removed. All indels were excluded from variant calling. Subsequently, the SNP matrix was filtered with BCFtools filter (v1.16) to discard sites with polymorphic probabilities under 99%; or genotypes with average supporting read depth less than 10*×*. Vcf2phylip (revision 0eb1b80) (https://github.com/edgardomortiz/vcf2phylip/tree/v2.0) was executed to create multi-sequence alignments from the filtered VCF file. For all species other than *S. cerevisiae*, maximum likelihood (ML) phylogenetic analysis was performed with RAxML (v8.2.12, “-f a -x 23333 -p 2333 –no-bfgs”) (Stamatakis, 2014) applying GTRGAMMA model and 100 times of bootstrap resampling. The host species trees were mid-point rooted and visualized in R (v4.2.3) using packages phytools (v1.5_1) (Revell, 2012) and ggtree (v3.6.0) (Yu *et al*., 2017), respectively. For *S. cerevisiae*, we followed the workflow in Peter *et al*. (2018): “snpgdsVCF2GDS” and “snpgdsDiss” from package SNPRelate (v1.32.0) (Zheng *et al*., 2012) were used to create the distance matrix from SNP data, and then function “bionj” from ape (v5.7_1) (Paradis *et al*., 2004) was used to reconstruct the neighbor joining (NJ) tree.

### Annotation and sequence analysis of full-length elements from whole genome assemblies

Ty elements were annotated in WGAs using a RepeatMasker-based pipeline previously described in Czaja *et al*. (2020) updated to use RepeatMasker v4.0.9. Three *S. cerevisiae* Ty4 elements with secondary FLE insertions from other Ty families were excluded from the final dataset to prevent multi-sequence alignment artifacts. *De novo* LTR element prediction was performed using LTRharvest (“-seed 100 -minlenltr 100 -maxlenltr 1000 -mindistltr 1500 -maxdistltr 15000 - similar 80.0 -xdrop 5 -mat 2 -mis -2 -ins -3 -del -3 -mintsd 5 -maxtsd 5 -motif tgca -motifmis 0 -vic 60 -overlaps best”) followed by LTRdigest (PFAM models: PF00078, PF00665, PF01021, PF03732, PF07727, PF12384, PF13976) in GenomeTools 1.6.1 (Ellinghaus *et al*., 2008; Steinbiss *et al*., 2009; Gremme *et al*., 2013; Mistry *et al*., 2021). Multi-sequence alignments of annotated FLEs were generated using MAFFT (v7.508) with default parameters (Katoh and Standley, 2013). Sub-regions of FLEs in alignments were identified by aligning the “Tsu4p_nw” sequence from a public database of annotated canonical yeast transposons (https://github.com/bergmanlab/yeast-transposons) with the FLE dataset, then selecting sub-regions with seqkit subseq (v0.16.1) (Shen *et al*., 2016). ML phylogenetic trees were reconstructed using RAxML (v8.2.12, “-f a -x 23333-p 2333 –no-bfgs”) (Stamatakis, 2014) with GTRGAMMA model and 100 bootstrap replicates. Phylogenetic network analysis was performed with SplitsTree4 (v4.15.1) (Huson and Bryant, 2006) applying the “Uncorrect_P” model and “NeighborNet” method. Pairwise sequence divergence was calculated based on Kimura’s 2-parameter substitution model with 50-bp sliding window size and 10-bp step size with R package spider (GitHub revision e93c5b4) (Brown *et al*., 2012) and phangorn (v2.11.1) (Schliep, 2011) in R (v4.2.3).

### Phylogenetic analysis of Tsu4 strain-specific consensus sequences

Strain-specific consensus sequences were generated with BCFtools (v1.16, “bcftools mpileup -a ‘FORMAT/DP’ -Q 20 -q 20”; “bcftools call -f GQ,GP -mv –skip-variants indels; bcftools consensus”) (Li, 2011) using BAM files previously generated by McClintock 2 (Chen *et al*., 2023). The percentage of bases supported by mapped reads (i.e., breadth) was calculated with BEDtools (v2.30.0, “bedtools genomecov - d -split”) (Quinlan and Hall, 2010). To avoid generating consensus sequences that are biased towards the Tsu4 reference sequence, samples with normalized Tsu4 depth less than 0.75 (estimated by McClintock coverage module) or breadth less than 0.9 (estimated by BEDtools genomecov) were removed from consensus sequence analysis. A multi-sequence alignment of strain-specific consensus sequences was generated using MAFFT (v7.508) (Katoh and Standley, 2013) with default parameters. ML phylogenetic analysis was performed using RAxML (v8.2.12, “-f a -x 23333 -p 2333 –no-bfgs”) applying GTRGAMMA model and 100 bootstrap replicates (Stamatakis, 2014). The ML tree was mid-point rooted using R package phytools (v1.2_0) (Revell, 2012) and then visualized using ggtree (v3.6.0) (Yu *et al*., 2017).

## Acknowledgments

We thank members of the Bergman and Garfinkel Labs for helpful discussion and comments during the project; the University of Georgia Genomics and Bioinformatics Core Facility (RRID:SCR_010994) for assistance with DNA extraction, PacBio library preparation and sequencing; and the Georgia Advanced Computing Resource Center for technical supporting and computational resources. This work was funded by the University of Georgia Research Foundation (CMB) and NIH grant R01GM124216 (DJG and CMB).

## Author Contributions

C.M.B. designed the experiments and computational analyses with input from J.C. and D.J.G. D.J.G. performed the cell culture experiments. J.C. developed computational pipelines for analysis of unassembled short read data, whole genome assembly, and phylogenetic analyses. C.M.B. developed computational pipelines for annotation of whole-genome assemblies. J.C. and C.M.B analyzed the data.

J.C. and C.M.B. drafted the manuscript with contributions from D.J.G. All authors revised and approved the final manuscript.

