## Supplemental Tables and Figures for "Horizontal transfer and recombination fuel Ty4 retrotransposon evolution in *Saccharomyces*"

### Supplementary Tables

Table S1: **Ty4 and Tsu4 subfamily content in whole genome assemblies of multiple *Saccharomyces* species.** Data for Ty4 and Tsu4 subfamily content for *S. cerevisiae* assemblies from [1] and assembly quality metadata for all assemblies can be found in Additional File 2. Five truncated Tsu4 copies in *S. kudriavzevii* IFO1802 (denoted with an asterisk) are divergent FLEs from a new subfamily in the Ty4 family.

| **Species** | **Strain** | **Ref.** | **Population** | **Strategy** | **# Ty4 full** | **# Ty4 trun.** | **# Ty4 solo** | **# Tsu4 full** | **# Tsu4 trun.** | **# Tsu4 solo** |
|---|---|---|---|---|---|---|---|---|---|---|
| *S. cerevisiae* | 245 | [2] | Mosaic | Illumina | 0 | 0 | 12 | 1 | 0 | 3 |
| *S. cerevisiae* | AFQ | [3] | Mosaic | Illumina | 0 | 10 | 24 | 1 | 0 | 1 |
| *S. cerevisiae* | CDM | [3] | Mosaic | Illumina | 0 | 7 | 32 | 0 | 1 | 3 |
| *S. cerevisiae* | CQS | [1] | French Guiana Human | ONT | 1 | 1 | 73 | 9 | 0 | 37 |
| *S. cerevisiae* | DBVPG6044 | [4] | West African | PacBio | 0 | 0 | 18 | 0 | 0 | 0 |
| *S. cerevisiae* | DBVPG6765 | [4] | Wine European | PacBio | 0 | 0 | 5 | 0 | 0 | 0 |
| *S. cerevisiae* | S288C | [4] | Laboratory | PacBio | 3 | 0 | 14 | 0 | 0 | 0 |
| *S. cerevisiae* | SK1 | [4] | Mosaic | PacBio | 0 | 0 | 14 | 0 | 0 | 0 |
| *S. cerevisiae* | UWOPS03-461.4 | [4] | Malaysian | PacBio | 0 | 0 | 60 | 0 | 0 | 0 |
| *S. cerevisiae* | Y12 | [4] | Asian fermentation | PacBio | 4 | 1 | 14 | 0 | 0 | 0 |
| *S. cerevisiae* | YPS128 | [4] | North American | PacBio | 4 | 0 | 13 | 0 | 0 | 0 |
| *S. paradoxus* | DG1768 | [5] | SpB | PacBio | 0 | 1 | 48 | 0 | 0 | 17 |
| *S. paradoxus* | MSH-604 | [6] | SpB | ONT | 0 | 1 | 46 | 4 | 0 | 19 |
| *S. paradoxus* | UFRJ50816 | [4] | SpB (S. America) | PacBio | 0 | 1 | 45 | 22 | 2 | 105 |
| *S. paradoxus* | YPS138 | [4] | SpB | PacBio | 0 | 0 | 49 | 1 | 0 | 18 |
| *S. paradoxus* | R23 | [6] | SpD | ONT | 0 | 1 | 51 | 4 | 0 | 29 |
| *S. paradoxus* | WX20 | [6] | SpD | ONT | 0 | 1 | 49 | 4 | 0 | 26 |
| *S. paradoxus* | LL2011_012 | [6] | SpC | ONT | 0 | 1 | 47 | 1 | 0 | 26 |
| *S. paradoxus* | LL2012_016 | [6] | SpC* | ONT | 0 | 1 | 46 | 1 | 1 | 33 |
| *S. paradoxus* | UWOPS91-917.1 | [4] | Hawaii | PacBio | 0 | 0 | 40 | 1 | 1 | 88 |
| *S. paradoxus* | LL2012_001 | [6] | SpA | ONT | 0 | 2 | 45 | 0 | 0 | 0 |
| *S. paradoxus* | CBS432 | [4] | EU | PacBio | 0 | 2 | 46 | 0 | 0 | 0 |
| *S. paradoxus* | N44 | [4] | FE | PacBio | 3 | 3 | 106 | 0 | 0 | 0 |
| *S. mikatae* | IFO 1815 | This study | Asia A | PacBio | 0 | 2 | 5 | 15 | 11 | 207 |
| *S. mikatae* | NBRC 10994 | This study | Asia C | PacBio | 0 | 1 | 10 | 26 | 8 | 234 |
| *S. jurei* | NCYC 3947 | [7] | Europe | PacBio | 0 | 0 | 7 | 4 | 2 | 137 |
| *S. jurei* | NCYC 3962 | [7] | Europe | PacBio | 0 | 0 | 6 | 3 | 2 | 138 |
| *S. kudriavzevii* | CR85 | [8] | Europe | ONT | 0 | 2 | 25 | 0 | 0 | 19 |
| *S. kudriavzevii* | ZP591 | [9] | Europe | PacBio | 0 | 2 | 22 | 0 | 0 | 18 |
| *S. kudriavzevii* | IFO1802 | [9] | Asia A | PacBio | 0 | 0 | 40 | 0 | 8* | 19 |
| *S. arboricola* | H-6 | [10] | Asia | 454 | 0 | 0 | 1 | 0 | 1 | 20 |
| *S. arboricola* | ZP960 | [11] | Oceania | Illumina | 0 | 0 | 0 | 0 | 1 | 26 |
| *S. uvarum* | CBS7001 | [12] | Holarctic | PacBio | 0 | 0 | 0 | 9 | 7 | 71 |
| *S. uvarum* | ZP964 | [9] | Australasia | PacBio | 0 | 0 | 0 | 0 | 6 | 74 |
| *S. eubayanus* | CBS12357 | [13] | Patagonia B | ONT | 0 | 0 | 1 | 0 | 6 | 98 |
| *S. eubayanus* | CL216.1 | [14] | Patagonia B | ONT | 0 | 0 | 1 | 0 | 7 | 100 |
| *S. eubayanus* | CL450.1 | [14] | Patagonia B | ONT | 0 | 0 | 1 | 4 | 10 | 107 |
| *S. eubayanus* | CDFM21L.1 | [15] | Holarctic | ONT | 0 | 0 | 1 | 14 | 4 | 160 |

Table S2: **Statistics for *de novo* whole genome assemblies of *S. mikatae* strains IFO 1815 and NBRC 10994 generated in this study.**

| | **IFO 1815** | **NBRC 10994** |
|---|---|---|
| # Contigs | 22 | 25 |
| Total Length (bp) | 12,243,664 | 12,361,694 |
| Contig N50 (bp) | 778,614 | 771,552 |
| L50 | 7 | 7 |
| GC content (%) | 37.8 | 37.99 |
| # tRNAs | 285 | 274 |
| Complete BUSCOs (%) | 97.19 | 97.23 |
| Data availability | JARBHO000000000 | JARBHP000000000 |

#### Supplementary Figures

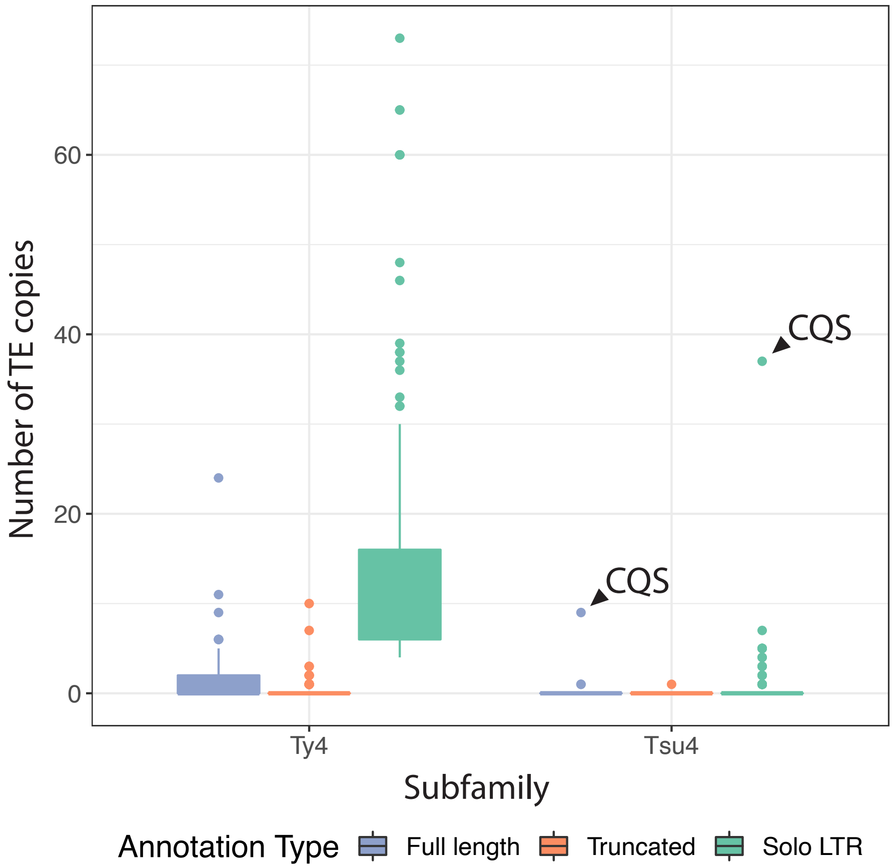


Figure S1: **Ty4/Tsu4 content in *S. cerevisiae* whole genome assemblies.** Shown are the box plots for numbers of Ty4 or Tsu4 copies annotated in each of 183 *S. cerevisiae* whole genome assemblies. Blue, red, and green boxes indicate FLEs, truncated elements, and solo LTRs, respectively. Colored boxes show interquartile ranges (IQR), whiskers show values 1.5×IQR of the upper or lower quartiles, and the dots indicate outliers that beyond 1.5×IQR. Outliers for Tsu4 elements in strain CQS are annotated.

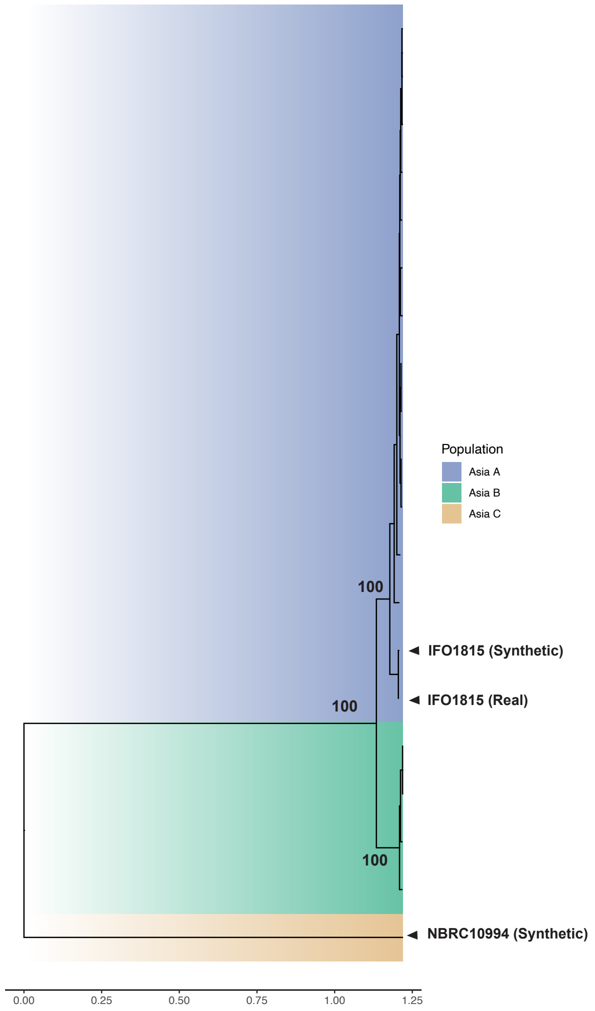


Figure S2: **Phylogenetic placement of IFO1815 and NBRC 10994 in *S. mikatae*.** ML tree based of *S. mikatae* strains based on empirical short-read WGS datasets from *S. mikatae* [11] plus short reads simulated from new IFO1815 and NBRC 10994 WGAs reported in this study. Clustering of IFO1815 samples based empirical [11] and simulated (this study) short read data confirms our simulation approach gives valid taxonomic placement, and reveals that NBRC 10994 represents a new *S. mikatae* clade (Asia C).

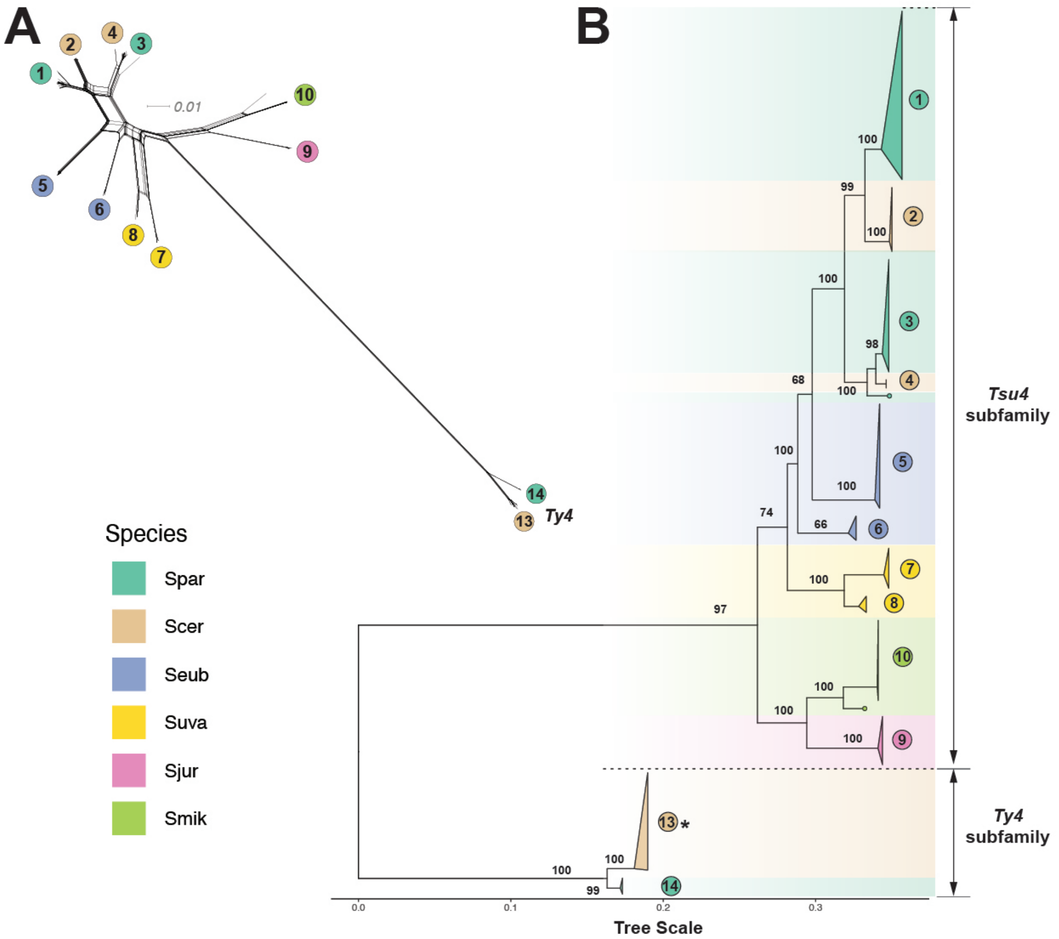


Figure S3: **Phylogenetic network and tree of FLEs from the Ty4 family in *Saccharomyces* excluding recombinant clades 11 and 12.** (A) Phylogenetic network for internal coding regions of Ty4/Tsu4 FLEs based on the NeighborNet algorithm. To simplify visualization, this network only includes Ty4 subfamily FLEs from WGAs reported in [4]. Lineages in the network are labeled according to monophyletic groups identified in Panel (B). Note that the signal for recombination between the Ty4 and Tsu4 subfamilies seen in Figure 3 is eliminated by exclusion of clades 11 and 12. (B) Midpoint rooted ML phylogeny of internal coding regions from Ty4/Tsu4 FLEs. Bootstrap support based on 100 replicates is shown for major nodes. The scale bar for branch lengths is in units of substitutions per site. All monophyletic groups are collapsed as triangles. Two singleton Tsu4 elements (f267 from Hawaiian *S. paradoxus* strain UWOPS91-917.1 and f256 from *S. mikatae* strain NBRC 10994) are denoted as dots at tips. Triangles, tip dots, and ranges are colored for each species. Vertical heights of triangles are proportional to the number of taxa. Horizontal widths of triangles are equal to the maximum branch length within the clade. Note that the monophyletic clade for the Ty4 subfamily from *S. cerevisiae* (annotated with an asterisk) is re-scaled to 5% of the real sample size both horizontally and vertically, due to the large number of Ty4 sequences (n=273) in *S. cerevisiae* genomes.

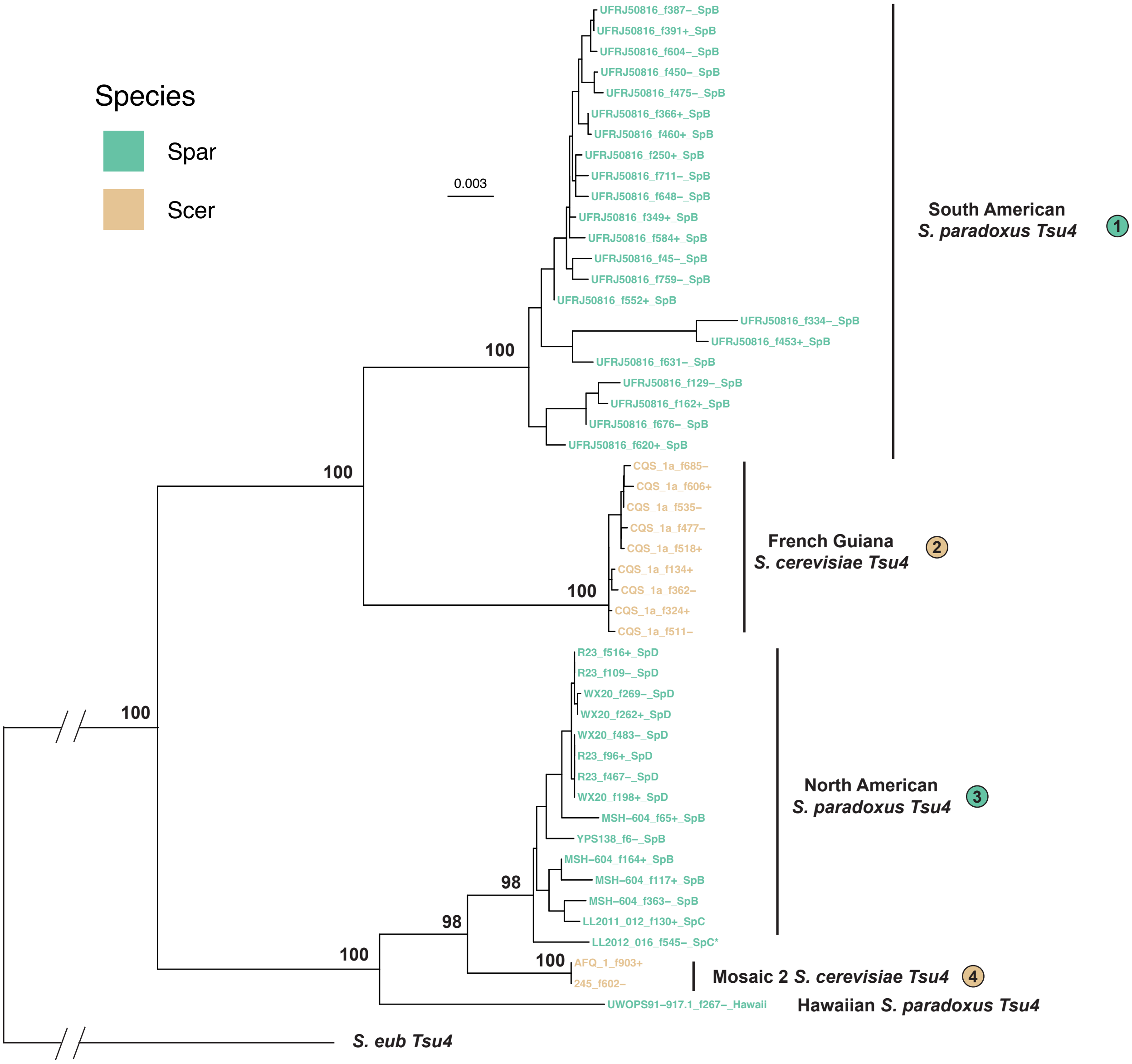


Figure S4: **Annotated phylogeny of Ty4/Tsu4 FLEs in *S. paradoxus* and *S. cerevisiae* genomes.** The phylogeny shown here is a rescaled, taxon-labeled sub-tree from Figure 3B, only showing the Tsu4 groups from *S. paradoxus* and *S. cerevisiae*. Bootstrap support based on 100 replicates, clade numbers are the same as in Figure 3B. The tree scale bar for branch lengths is in units of substitutions per site. For *S. paradoxus* clades, the geographic source is annotated, and taxon labels consist of strain identifier, FLE identifier and host lineage. For *S. cerevisiae* clades, the host lineage is annotated, and taxon labels consist of strain identifier and FLE identifier.

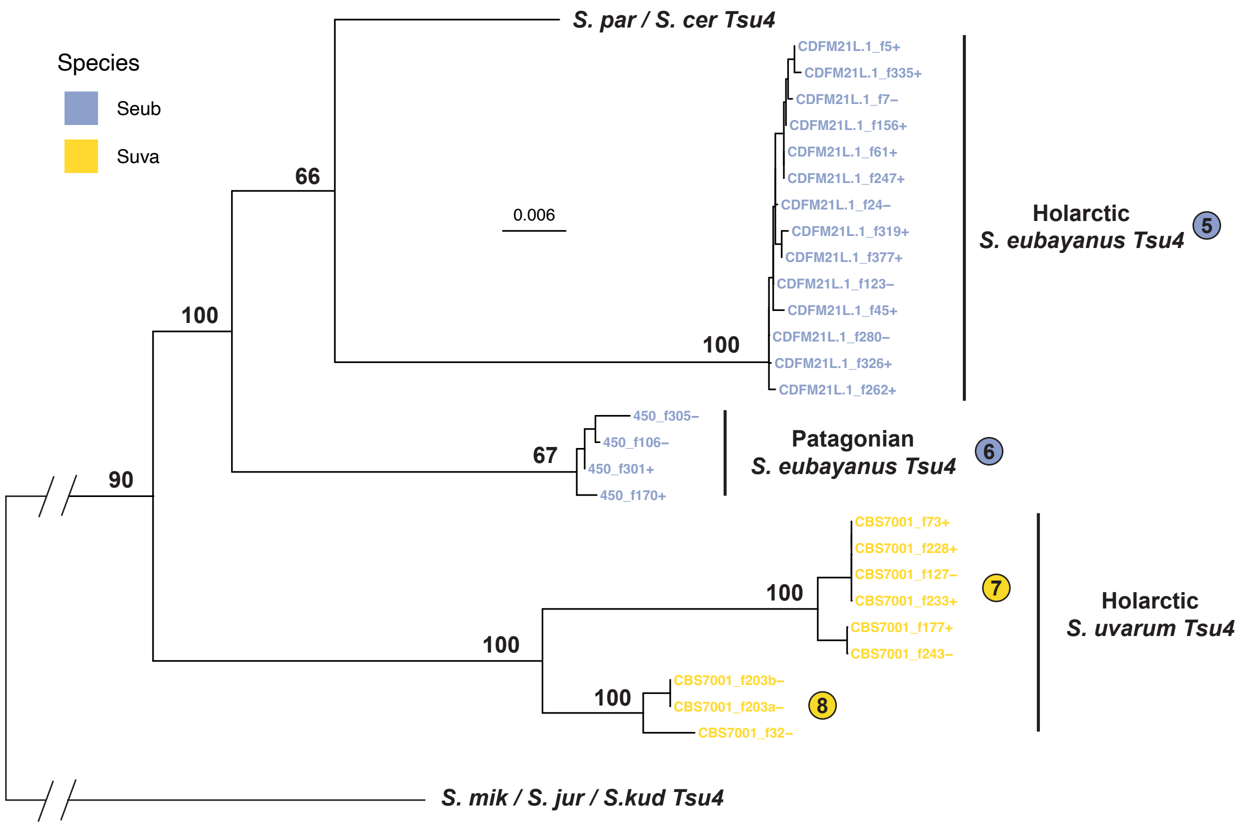


Figure S5: **Annotated phylogeny of Ty4/Tsu4 FLEs in *S. uvarum* and *S. eubayanus* genomes.** The phylogeny shown here is a rescaled, taxon-labeled sub-tree from Figure 3B, only showing the Tsu4 groups from *S. uvarum* and *S. eubayanus*. Bootstrap support based on 100 replicates, clade numbers are the same as in Figure 3B. The tree scale bar for branch lengths is in units of substitutions per site. The host lineage is annotated and taxon labels consist of strain identifier and FLE identifier for all clades.

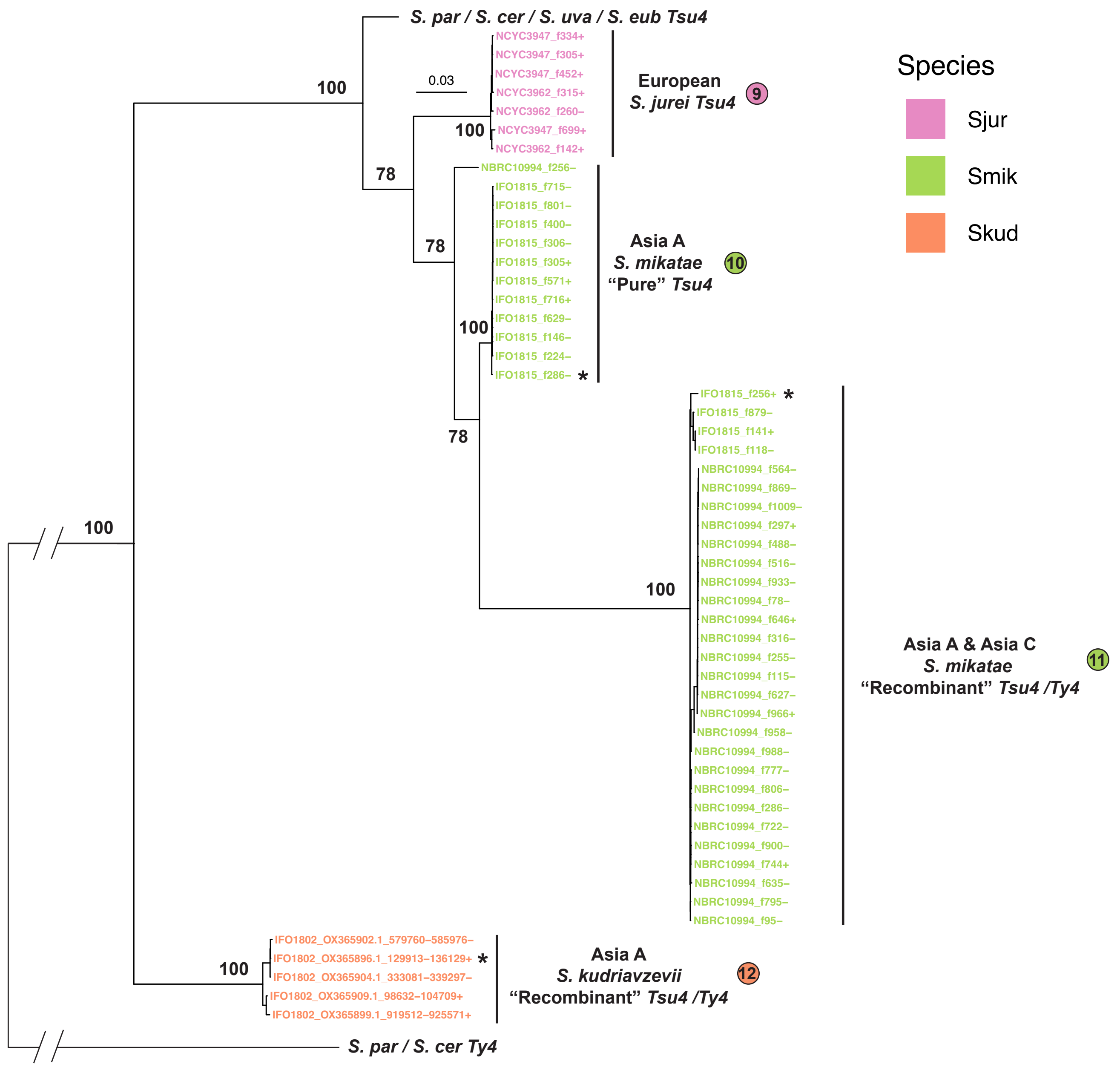


Figure S6: **Annotated phylogeny of Ty4/Tsu4 FLEs in *S. mikatae*, *S. jurei* and *S. kudriavzevii* genomes.** The phylogeny shown here is a rescaled, taxon-labeled sub-tree from Figure 3B, only showing the Tsu4 clades from *S. mikatae*, *S. jurei* and *S. kudriavzevii*. Bootstrap support based on 100 replicates, clade numbers are the same as in Figure 3. The tree scale bar for branch lengths is in units of substitutions per site. Taxon labels consist of strain identifier and FLE identifier for all clades. Representative FLEs from *S. mikatae* and *S. kudriavzevii* recombinant clades 11 and 12 selected for sliding window divergence analysis are labeled with asterisks.

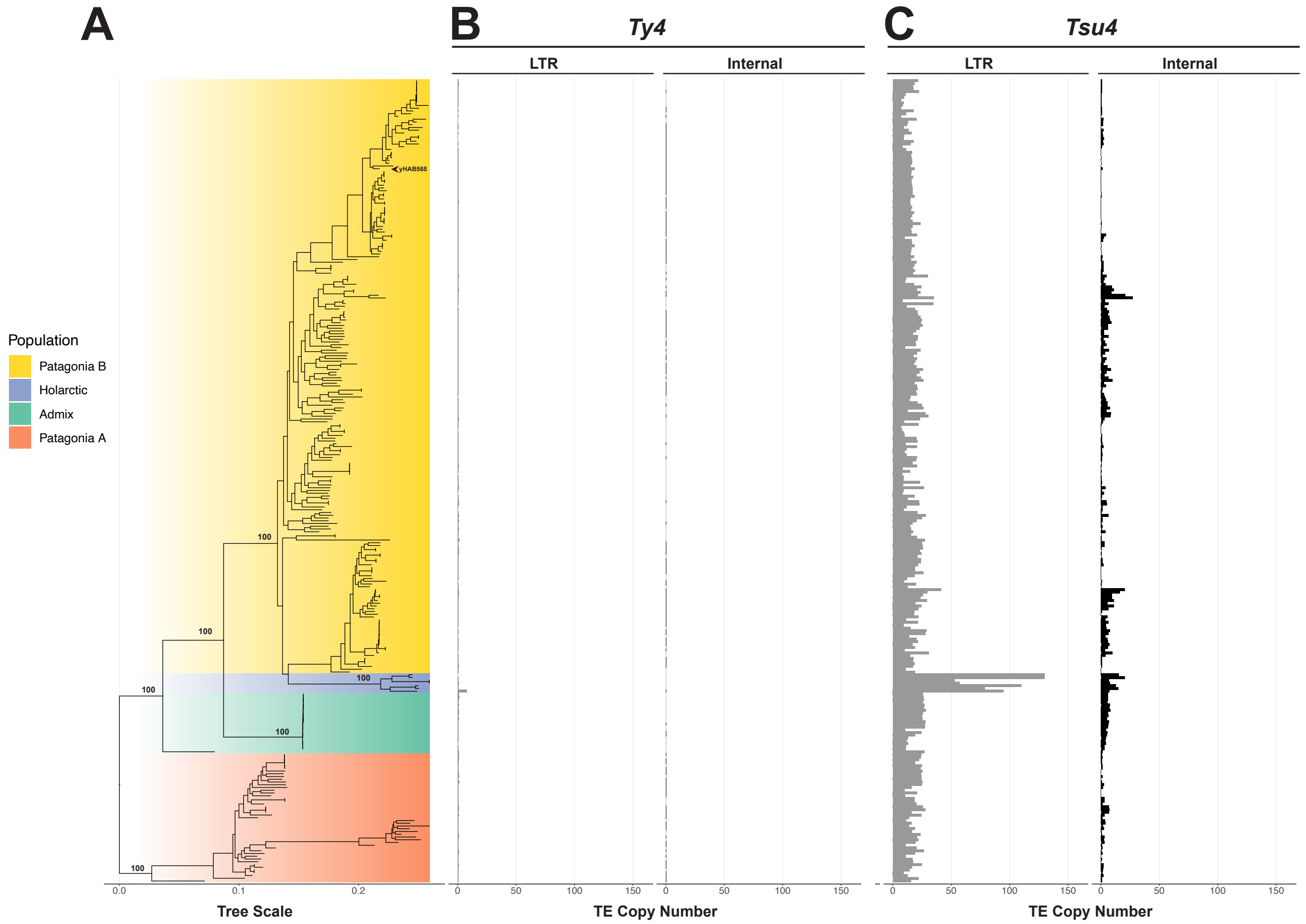


Figure S7: **Host phylogeny of *S. eubayanus* annotated with Ty4/Tsu4 copy number estimates.** Shown are the copy number estimates for Ty4 (B) and Tsu4 (C) subfamilies from worldwide *S. eubayanus* lineages. The ML tree is reconstructed using 319,865 genome-wide SNPs from 292 *S. eubayanus* strains and midpoint rooted. Plotting details are identical as described in Figure 1. Major lineages are annotated according to previously-reported population structure [16]. *S. eubayanus* strain yHAB565 whose strain-specific Tsu4 consensus sequence clusters with *S. uvarum* is indicated with an arrowhead in the Patagonia B lineage.

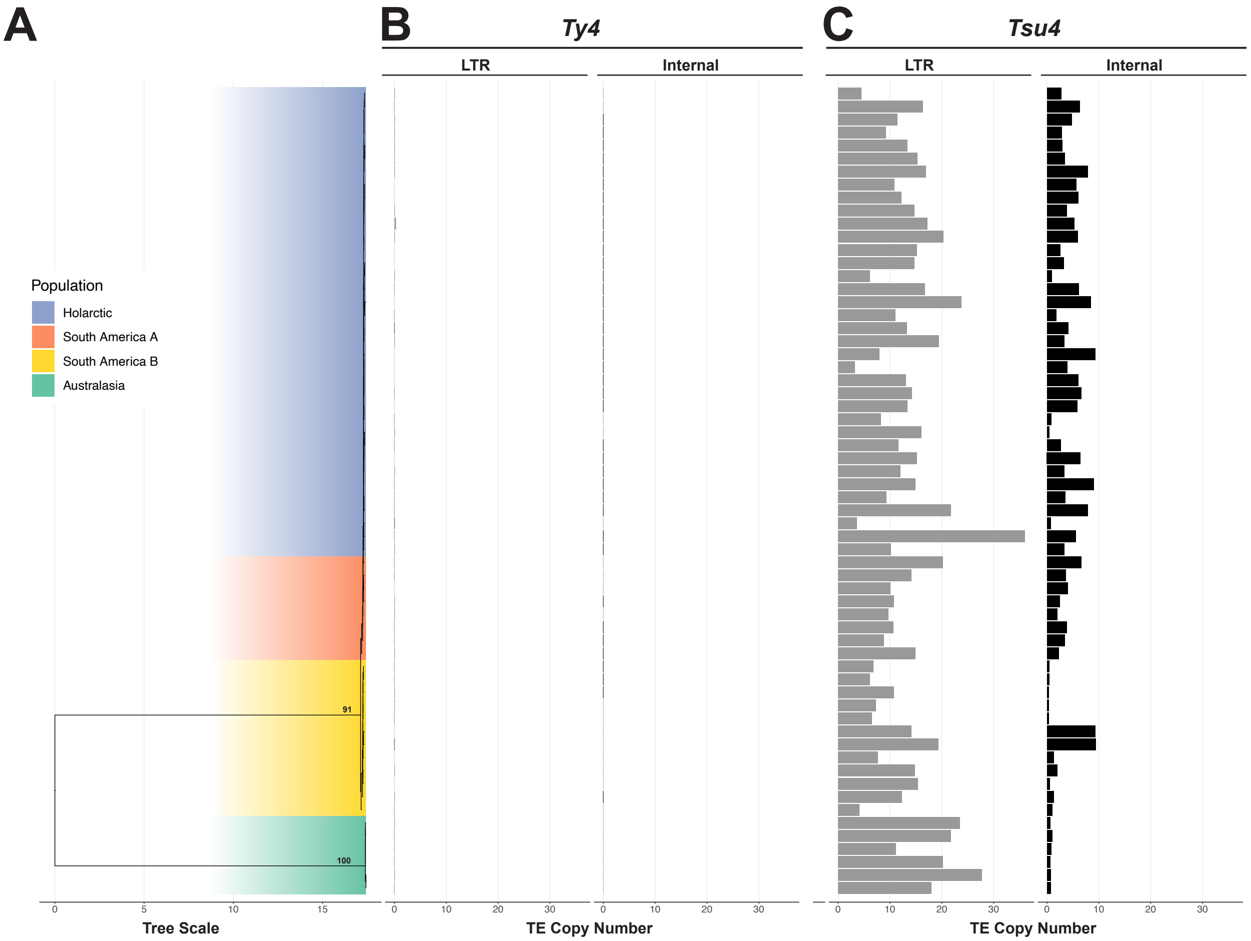


Figure S8: **Host phylogeny of *S. uvarum* annotated with Ty4/Tsu4 copy number estimates.** Shown are the copy number estimates for Ty4 (B) and Tsu4 (C) subfamilies from worldwide *S. uvarum* lineages. The ML tree is reconstructed using 253,252 genome-wide SNPs from 62 *S. uvarum* strains and midpoint rooted. Plotting details are identical as described in Figure 1. Major lineages are annotated according to previously-reported population structure [17].

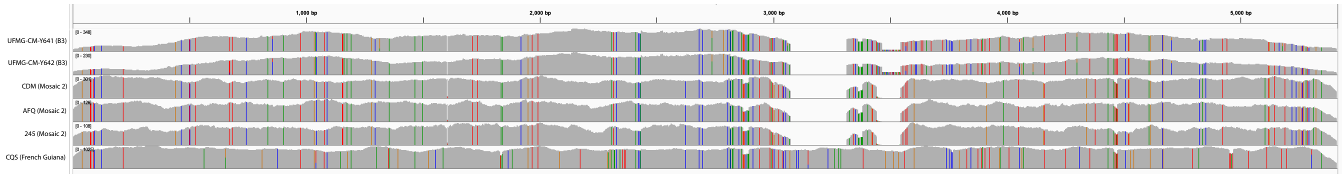


Figure S9: **Patterns of single nucleotide variation in *S. cerevisiae* strains with Tsu4 internal regions.** Integrated Genomics Viewer [18] screenshot of single nucleotide variation relative to the *S. paradoxus* Tsu4 reference seqeuence for six *S. cerevisiae* strains with Tsu4 internal region copy number >0.5. Variant profiles cluster into three groups: one containing the two B3 lineage strains UFMG-CM-Y641 and UFMG-CM-Y642; one containing the three Mosaic 2 lineage strains CDM, AFQ, and 245; and one containing the French Guiana lineage strain CQS.

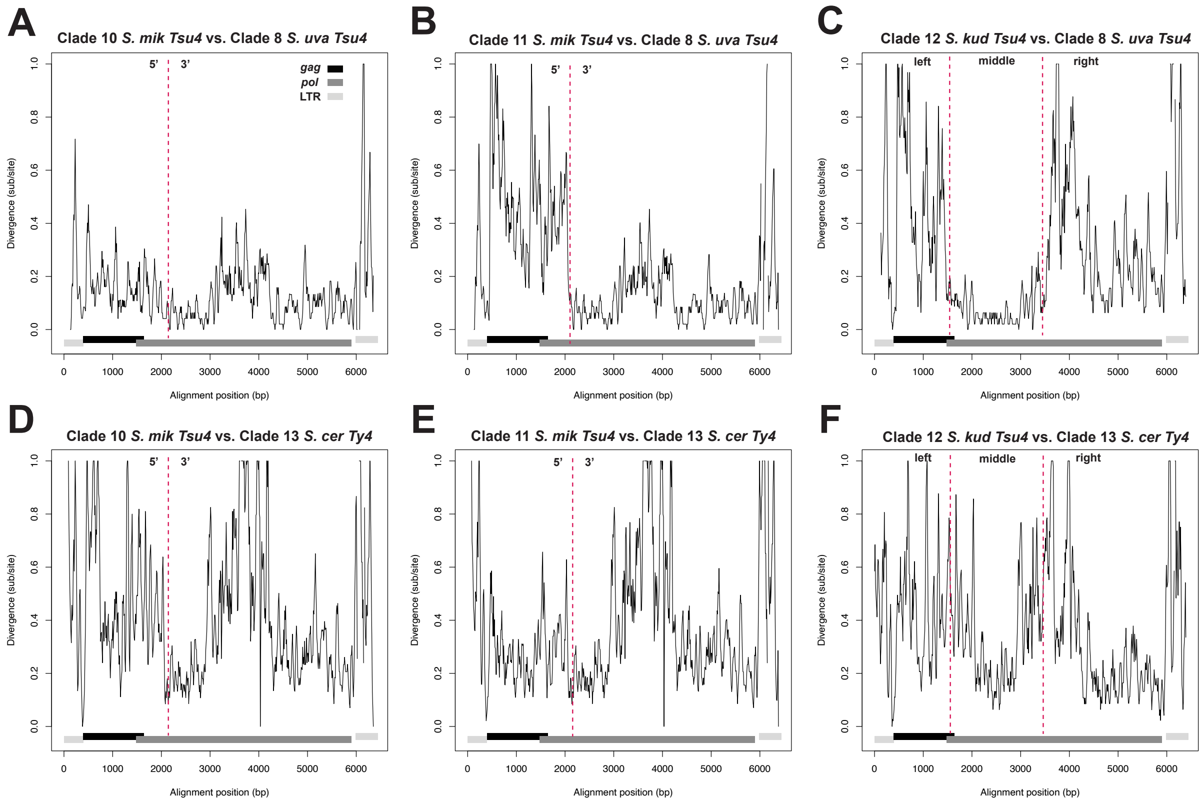


Figure S10: **Sequence divergence between recombinant and pure Tsu4 from *S. mikatae* versus *S. uvarum* Tsu4 and *S. cerevisiae* Ty4.** Shown are sliding window analysis of pairwise sequence divergence between (A) "Recombinant" Tsu4 in *S. mikatae* vs. Tsu4 in *S. uvarum*; (B) "Pure" Tsu4 in *S. mikatae* vs. Tsu4 in *S. uvarum*; (C) "Recombinant" Tsu4 in *S. mikatae* vs. Ty4 in *S. cerevisiae*; (D) "Pure" Tsu4 in *S. mikatae* vs. Ty4 in *S. cerevisiae*. Structure of a Ty4 FLE is annotated at the bottom of each panel in colored rectangles (*Gag* in black, *Pol* in darker gray, and LTRs in lighter gray). The dashed red line in panel (A) indicates the boundary of 5' and 3' internal region which is later used for partitioning the Ty4/Tsu4 phylogeny in Figure 5. Elements used in this analysis include IFO1815_f256 for "Recombinant" Tsu4 in *S. mikatae*, IFO1815_f286 for "Pure" Tsu4 in *S. mikatae*, CBS7001_f32 for Tsu4 in *S. uvarum*, YPS128_f49 for Ty4 in *S. cerevisiae*. IFO 1815 assembly was generated in this study. CBS 7001 and YPS128 assemblies were previous published in Chen *et al.* [12] and Yue *et al.*[4], respectively. Divergence measured in substitutions per site was calculated using a Kimura 2-parameter model in overlapping 50 bp windows with a 10 bp step size.
